# Benchmarking residue-resolution protein coarse-grained models for simulations of biomolecular condensates

**DOI:** 10.1101/2024.08.28.610132

**Authors:** Alejandro Feito, Ignacio Sanchez-Burgos, Ignacio Tejero, Eduardo Sanz, Antonio Rey, Rosana Collepardo-Guevara, Andres R. Tejedor, Jorge R. Espinosa

## Abstract

Intracellular liquid–liquid phase separation (LLPS) of proteins and nucleic acids is a fundamental mechanism by which cells compartmentalize their components and perform essential biological functions. Molecular simulations play a crucial role in providing microscopic insights into the physicochemical processes driving this phenomenon. In this study, we systematically compare six state-of-the-art sequence-dependent, residue-resolution models to evaluate their performance in reproducing the phase behaviour and material properties of condensates formed by seven variants of the low-complexity domain (LCD) of the hnRNPA1 protein (A1-LCD)—a protein implicated in the pathological liquid-to-solid transition of stress granules. Specifically, we assess the HPS, HPS-cation–*π*, HPS-Urry, CALVADOS2, Mpipi, and Mpipi-Recharged models in their predictions of the condensate saturation concentration, critical solution temperature, and condensate viscosity for the A1-LCD variants. Our analyses demonstrate that, among the tested models, Mpipi, Mpipi-Recharged, and CALVADOS2 provide accurate descriptions of the critical solution temperatures and saturation concentrations for the various A1-LCD variants tested. Regarding the prediction of material properties for condensates of A1-LCD and its variants, Mpipi-Recharged stands out as the most reliable model. Overall, this study benchmarks a range of residue-resolution coarse-grained models for the study of the thermodynamic stability and material properties of condensates and establishes a direct link between their performance and the ranking of intermolecular interactions these models consider.

## I. AUTHOR SUMMARY

Molecular simulations have proved to be invaluable for gaining microscopic insights of the physicochemical processes underlying the formation of membraneless organelles—e.g., biomolecular condensates—in the cell. Computational efficiency and predictive accuracy are mandatory requirements for biophysical models aiming to elucidate the phase behaviour of condensates. In this study, we evaluate the performance of several state-of-the-art residue-resolution protein models in describing the phase diagram and material properties of condensates formed by different intrinsically disordered mutants of the low-complexity domain of hnRNPA1. This protein plays a key role in the stabilization and pathological solidification of the stress granules, a type of cytoplasmic condensates whose liquid-to-solid transition has been associated to the onset of multiple neurodegenerative dis-orders. Our results, besides serving as a powerful benchmark for different models to describe protein liquid-liquid phase-separation, further establish a direct relation between condensate phase behaviour and individual intermolecular interactions. We conclude that the central intermolecular interactions dictating the phase behaviour of the hnRNPA1 low-complexity domain—the key protein domain involved in phase-separation and fibrillization—are primarily cation-*π* interactions as arginine-tyrosine and arginine-phenylalanine, and *π*-*π* interactions mediated by tyrosine and phenylalanine contacts.

## II. INTRODUCTION

The formation of intracellular membraneless organelles (MLOs), known as biomolecular condensates, represents an essential mechanism enabling the spatio-temporal organisation and functional regulation of the cell material^1–3^. Biomolecular condensates contain a wide variety of biomolecules, including intrinsically disordered proteins (IDPs), multi-domain proteins, and DNA or RNA strands. These condensates are thought to form via spontaneous demixing of biomolecules by means of liquid-liquid phase separation (LLPS). Biomolecular condensates have been linked to diverse biological functions, such as cell signalling^4–6^, buffering protein concentrations^7,8^, compartmentalisation^7–10^, genome silencing^11,12^, or noise buffering^13–15^, among others^16,17^. Many naturally occurring phase-separating proteins—such as the widely studied *fused in sarcoma* (FUS)^18–20^, the TAR DNA-binding protein of 43kDa (TDP-43)^20–22^, or the heterogeneous nuclear ribonucleoprotein A1 (hn-RNPA1)^23,24^—are multi-domain proteins that contained intrinsically disordered regions (IDRs). IDRs confer multi-valency to such proteins, as they can establish multiple transient intermolecular interactions, which have been consistently shown to promote condensate formation via phase separation^25–28^. Some of the IDRs present in phase-separating proteins are characterized by having amino acid sequences of low complexity, and thus are termed low-complexity domains (LCD). Besides promoting condensate formation, LCDs can also trigger the progressive rigidification of liquid-like biomolecular condensates into solid-like states^29–31^. The liquid-to-solid transitions of condensates has been linked to the onset of several neurodegenerative pathologies such as amyotrophic lateral sclerosis (ALS)^32–34^, frontotemporal dementia (FTD)^35,36^, Alzheimer^37,38^, or even some types of cancer^17,39,40^. Thus, unravelling the conditions, factors, and interactions regulating protein self-assembly, and their subsequent potential liquid-to-solid transition into aberrant solid states represents an urgent challenge in cell biology^25,34^.

Uncovering the factors governing the phase behaviour of biomolecules represents a complex challenge, necessitating an integrated approach that combines experimental and computational methodologies^2,41,42^. Biomolecular modelling and simulations, in particular, play a pivotal role in unravelling the underlying mechanisms and parameters driving phase separation. These approaches offer detailed insights into the forces mediating interactions between biomolecules, enhancing our understanding of the processes that govern their assembly^41–43^. Biomolecular simulations can widely vary in resolution and performance of the models used, spanning from atomistic models^44–47^—where all atoms in the system are described explicitly but only a few proteins or protein segments can be studied—to coarsegrained models^48–51^—where groups of several atoms, or even whole proteins, are represented by a single particle and the interactions among particles are simplified to improve computational efficiency, allowing the investigation of systems with hundreds of molecules. Amongst these, so-called ‘sequence-dependent’ coarsegrained models, which have a resolution of one-bead per amino acid, have become the method of choice for probing the link between amino acid sequence and phase behaviour^41,52–61^. These models have been used to gain molecular mechanisms explaining the phase behaviour of many different proteins^62–68^, as well as the impact of other biomolecules—e.g. RNA, DNA and chromatin—in controlling condensate architecture, stability, or transport properties of condensates^47,69,70^. Through iterative refinement and validation against experimental benchmarks, these sequence-dependent coarse-grained models are continually advancing and provide valuable insights into the intricate relationships between protein sequence, structure, dynamics, and their collective phase behaviour^42,52^.

Experimental measurements are one of the fundamental baselines to guide the development and testing the performance of sequence-dependent models for studying protein phase behaviour^52,53^. Approaches for the parameterisation of sequence-dependent coarse-grained models include using experimentally-derived hydrophobicity scales^43,54^ machine-learning algorithms^41,55,57^, and combining bioinformatic analyses with atomistic simulations^53,61^. A common approach for testing the performance of sequence-dependent coarse-grained models has been comparison against *in vitro* measurements of singlemolecule radius of gyration (*R*_*g*_)^71–75^ of IDRs^41,56,59,76^.

Sequence-dependent coarse-grained models, like Mpipi^53^ and Mpipi-Recharged^61^, have been tested instead by comparing their predictions against experimental phase diagrams of protein solutions^77^. Therefore, experimental efforts providing temperature-dependent coexistence lines, single-protein radius of gyration, or viscosity values are enormously valuable, aside of their own interest, to assess the predictive capability of coarse-grained (CG) models^77,78^.

In this work, we benchmark the performance of six residue-resolution CG models by evaluating their predictions of the saturation concentration and temperature-vs-density phase diagrams for several hnRNPA1-LCD (referred to as A1-LCD hereafter) mutants, comparing them against *in vitro* experimental data^77,78^. The models under evaluation are: HPS^43^, HPS-cation– *π*^56^, HPS-Urry^54^, CALVADOS2^55^, Mpipi^79^, and Mpipi-Recharged^61^. Our analysis highlights the sensitivity of these coarse-grained models in capturing the effects of sequence modifications on the propensity of A1-LCD condensates to form. In addition, we test ability of the models to capture condensate viscosity. This is a particularly challenging benchmark, as condensate viscosity is not an explicit target property in the parametrization of any of these models. Nevertheless, we consider this a critical test since accurately capturing viscosity reflects the models’ capacity to balance the diverse biomolecular forces that stabilize condensates. Thus, how well the models predict condensate viscosity is directly linked to the quality of their parametrizations of amino acid pair interactions. After establishing the differences in the model predictions for the the experimental phase behaviour of A1-LCD^77,78^, we analyse the protein intermolecular contact maps and the most frequent residue–residue interactions within condensates for the studied mutants, with the aim to elucidate key differences among the various models. Overall, our findings contribute to the refinement of residue-resolution coarse-grained models and describe how changes in model parametrisations impact differently a range of biophysical properties of condensates.

## III. RESULTS

### A. Prediction of A1-LCD coexistence lines using different sequence-dependent models

The determination of the phase diagram establishes the thermodynamic conditions to investigate the formation and stability of biomolecular condensates^80,81^. Numerous experiments have consistently shown how aromatic residues represent key ‘stickers’ to sustain protein phase separation of prion-like domain proteins^23,77,78,82,83^. Therefore, to evaluate the performance of different models such as the HPS^52^, HPS-cation– *π*^56^, HPS-Urry^54^, CALVADOS2^55^, Mpipi^53^ and Mpipi-Recharged^61^ in reproducing condensate behaviour (see Section V A for technical details about the models), we consider the different mutants of A1-LCD reported in Ref.^78^ that display multiple variations in the type of aromatic residues along their sequence (see Table A in Supporting Information (SI) to visualize the studied sequences). These variants are the following: (1) WT+NLS which corresponds to the A1-LCD wild-type sequence, which features a nuclear localization signal^78^ and contains 8 tyrosines (Y) and 12 phenylalanines (F); (2) allF with 19F; (3) allY with 19Y; (4) allW with 19 tryptophans (W); (5) YtoW with 7W and 12F; (6) FtoW with 7Y and 12W; and (7) W^*−*^ with 13W. Importantly, the mutations in all variants are performed on the aromatic residues maintaining the patterning and just changing the chemical identity of the residues^78^.

We next compute the temperature-vs-density phase diagrams of the aforementioned sequences by performing Direct Coexistence (DC)^84,85^ molecular dynamics simulations of protein solutions (see Fig. 1A and Section V C for further methodological details). In the DC method, the two coexisting phases are simulated by preparing periodically replicated slabs of the two phases (the condensed and the diluted phase) in the same simulation box. Once the system is equilibrated, a density profile along the long axis of the box can be extracted to compute the density (or concentration) of the two coexisting phases after a production run of *∼* 1*μ*s (please see Supporting Movie A for a representative trajectory of our DC simulations). Importantly, we note that none of the six employed models includes explicit solvent, hence, effectively in our DC simulations the protein diluted phase corresponds to a vapour phase. In Fig. 1A, a typical snapshot of the simulation slab is shown together with the associated density profile below the critical temperature (Top panel), and above the critical point (Bottom panel), where a single phase is present.

**FIG. 1.**
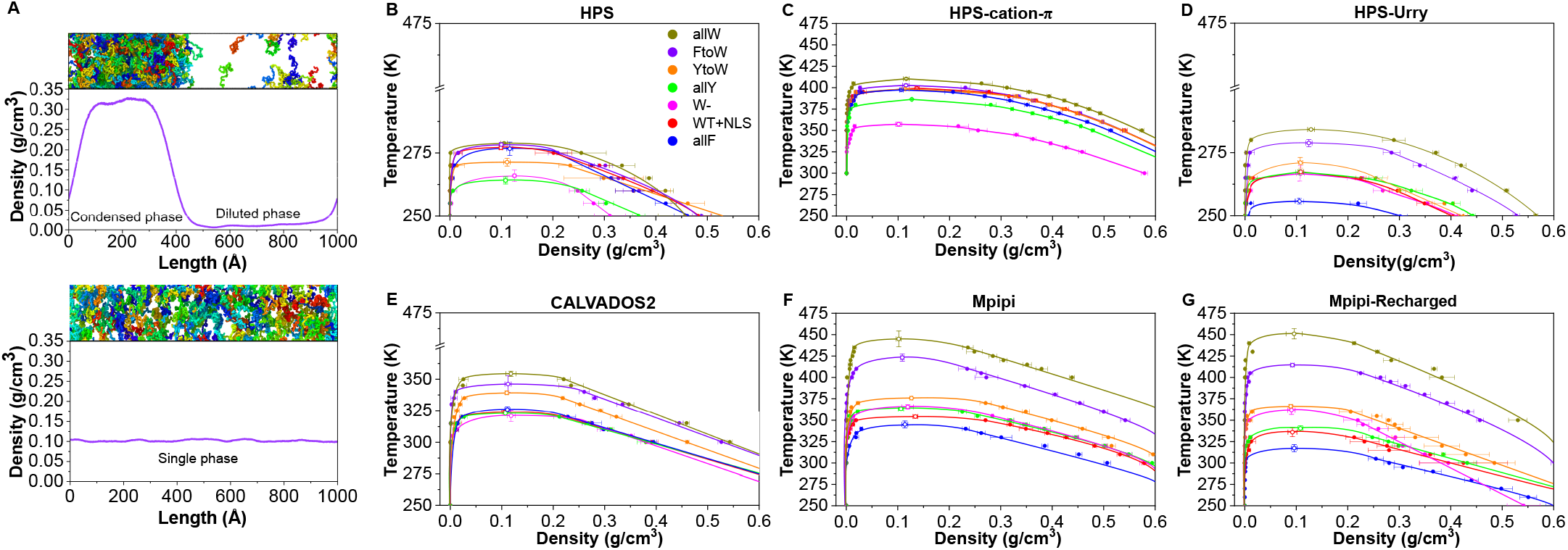
(**A**) Snapshots of a Direct Coexistence (DC) simulation used to calculate the phase diagram and critical temperature of A1-LCD (WT+NLS) protein, where each protein replica has a different colour. Top panel exhibits phase separation with a distinguishable condensed and diluted phase as depicted by the associated density profile. Bottom panel represents a system above the critical temperature where no phase separation occurs and a single phase is detected across the density profile. **B**-**G** Temperature–density phase diagrams of A1-LCD variants calculated via DC simulations employing the HPS (**B**), HPS-cation–*π* (**C**),HPS-Urry (**D**), CALVADOS2 (**E**), Mpipi (**F**) and Mpipi-Recharged (**G**) models. Critical temperatures are represented by empty circles while filled circles depict coexistence densities obtained from DC simulations. The lines serve as a guide to the eye. The colour coding is preserved throughout all the panels for all the variants as indicated in the legend of panel B.

The phase diagrams for the seven A1-LCD variants, computed using six different models, are shown in Fig. 1B-G. These diagrams highlight the varying sensitivity of the models in capturing the effects of specific amino acid mutations across the protein sequence. Among the models, Mpipi and Mpipi-Recharged (Fig.1F and G, respectively) display the largest variation in the critical temperature (T_*c*_), with T_*c*_ spanning more than 100 K from the lowest to the highest value across the variants. In contrast, the T_*c*_ values predicted by HPS and HPS-family models vary by only about 30 K between the lowest and highest values (Fig. 1B and D). Despite this, the HPS-Urry model (Fig. 1D) demonstrates greater sensitivity to amino acid mutations, as evidenced by the larger spacing in T_*c*_ values among the variants.

The HPS model^76^ represents amino acid pair interactions using a Lennard-Jones potential parametrized based on the Kapcha and Rossky (KR) hydrophobicity scale^86^, derived from the atomic partial charges of amino acids extracted from an all-atom force field. Specifically, the hydrophobicity values from the KR scale for each amino acid (*λ*_*i*_, where *i* is the amino acid) are used to define the interaction strength parameter (*ϵ*_*i*_ = *cλ*_*i*_, where *c* is a scaling constant) for the Lennard-Jones potential for each amino acid type. To determine the Lennard-Jones parameters for interactions between different amino acid types, the model uses the Lorentz-Berthelot mixing rules. The HPS model also incorporates salt-screened charge– charge interactions modelled through a Debye-Hückel potential. The HPS-cation–*π* model^56^ is a reparametrization of the HPS model that enhances the relative strength of all the cation–*π* pair interactions to better capture the experimentally observed phase separation propensity of DDX4 IDR variants^87^. More recently, the HPS-Urry model^54^ was developed to improve the predictive accuracy of the HPS family of models. This version replaces the KR hydrophobicity scale with the Urry hydrophobicity scale^88^, enabling improved predictions of the effects of R-to-K and Y-to-F mutations on the phase behaviour of protein solutions.

The CALVADOS family of models^41,55^, including the CALVADOS2 model tested here, builds on the foundation of the HPS model. However, instead of using a hydrophobicity scale to define the *λ*_*i*_ values, and subsequently the Lennard-Jones parameters, the CALVADOS models employ a Bayesian learning approach. In this approach, the *λ*_*i*_ values for all amino acids are optimized to achieve agreement with experimental single-molecule radii of gyration and paramagnetic relaxation enhancement (PRE) nuclear magnetic resonance (NMR) data across a broad range of intrinsically disordered regions (IDRs).

In contrast, the Mpipi model^53^, developed by our group, takes a fundamentally different approach by abandoning the Lorentz-Berthelot mixing rules and instead defining amino acid pair interactions in a pair-specific manner. Additionally, the model uses the Wang-Frenkel potential, which offers greater flexibility and computational efficiency compared to the Lennard-Jones potential. For charge–charge interactions, Mpipi employs a Debye-Hückel potential, similar to the HPS and CALVADOS families. However, we recently introduced the Mpipi-Recharged model^61^, which enhances the original Mpipi model by refining its description of electrostatic interactions. Specifically, Mpipi-Recharged replaces the pair-agnostic Debye-Hückel potential with a pair-specific, non-symmetric Yukawa potential. This parameterization is based on atomistic simulations of amino acid pairs in explicit water with ions, allowing for a more accurate depiction of charge effects. By abandoning mixing rules in the Wang-Frenkel potential (in both Mpipi and Mpipi-Recharged models) and incorporating amino acid-specific descriptions of charge interactions (in Mpipi-Recharged), the Mpipi family of models addresses critical limitations of standard mixing rules. These rules may fail to capture nuanced biomolecular interactions, such as the subtle balance between aromatic stacking, cation–*π* interactions, and charge–charge interactions. Treating pair interactions explicitly enables the Mpipi models to achieve excellent agreement with experimental data^53,61,68,89^. This pair-specific approach not only improves accuracy but also provides a more detailed understanding of the molecular forces driving phase behaviour^61^.

To provide additional context on the differences between the models and their resulting phase diagrams, Fig. A (in SI) presents the relative interaction strength maps for each model, encompassing both hydrophobic and electrostatic interactions between all amino acid pairs. These interaction maps reveal significant differences in how the models parametrize interactions for the aromatic residues Y, F, and W. In the HPS and HPS-cation–*π* models, the strength of interactions between aromatic residues and the rest of the amino acids are relatively homogeneous. In contrast, Mpipi, Mpipi-Recharged, CALVADOS2, and HPS-Urry display more pronounced differences in their energy scales, with W *>* Y *>* F in terms of interaction strength. The variations in the energy scales of the models and the relative strength of interactions among amino acid pairs strongly influence how sensitive each model is to specific (aromatic) sequence mutations and their impact on the encoded phase behaviour. Models with greater differences in interaction strengths for aromatic residues tend to show enhanced sensitivity to mutations, as reflected in their broader T_*c*_ ranges and the corresponding phase diagrams.

The temperature-vs-concentration phase diagram of three of these sequences (WT+NLS, allY, and allF) has been recently measured *in vitro*^77,78^. By fitting critical solution temperatures from the experimental data, we now establish a direct comparison between modelling and experiments so we can assess the ability of the models to predict A1-LCD phase behaviour. Accordingly, the phase diagrams in Fig. 2 are plotted as a function of the protein concentration (and in logarithmic scale) to evaluate their predictive capability against experiments in the diluted phase. Determining the protein concentration in the dilute phase—specifically, the saturation concentration— poses significant challenges for Direct Coexistence simulations, particularly at low temperatures, due to the limited sampling inherent to these simulations^68^.

**FIG. 2.**
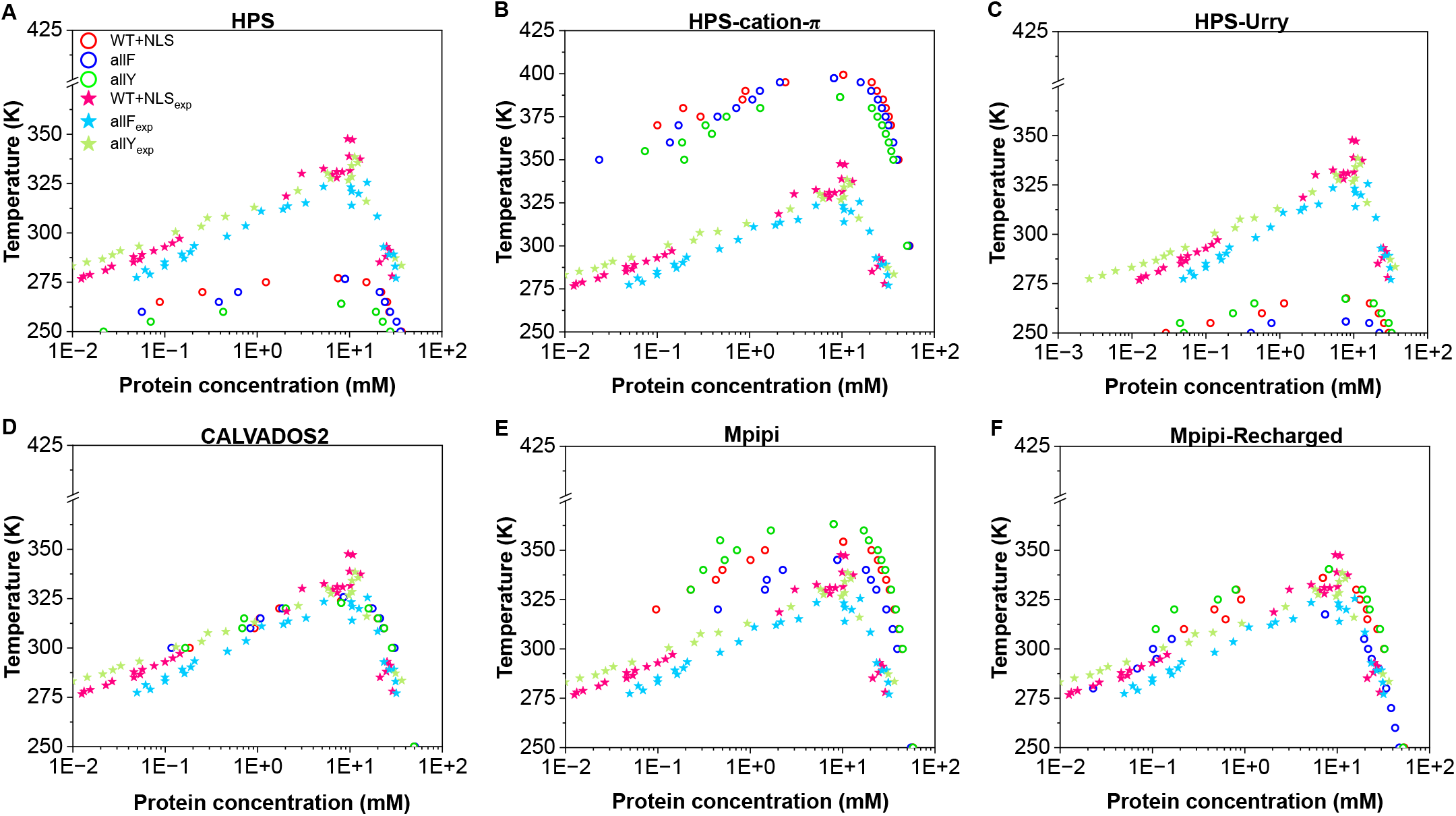
Experimental (solid stars) *vs*. simulated (empty circles) phase diagrams for the WT+NLS, allF, and allY variants of A1-LCD using different models: HPS (**A**), HPS-cation–*π* (**B**), HPS-Urry (**C**), CALVADOS2 (**D**), Mpipi (**E**), and Mpipi-Recharged (**F**).

As illustrated in Fig. 2A, the HPS model^76^ consistently underestimates the critical solution temperatures for the three A1-LCD variants tested. When comparing the predicted critical temperatures from simulations (orange symbols in Fig. 3) with the corresponding experimental values (derived by us from coexistence concentrations reported in Ref.^77^), the trend line deviates significantly from the perfect-fit reference (black line with a slope of 1 and intercept of 0; Fig. 3). This suggests that the HPS model does not effectively capture the impact of aromatic amino acid mutations on the critical solution temperatures of the A1-LCD system. The discrepancy is likely due to the parametrization of this early version of the HPS model, which relies on the KR hydrophobicity scale^76,86^. In the KS scale, F is ranked as more hydrophobic than W, and W is ranked as more hydrophobic than Y, potentially misrepresenting the relative contributions that these aromatic residues have in the phase separation of these systems.

**FIG. 3.**
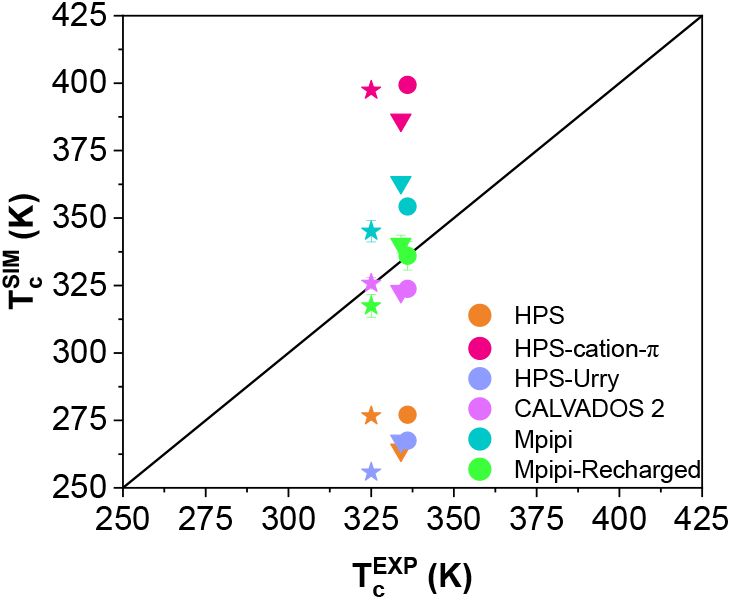
Comparison of the critical temperatures of WT+NLS (circles), allF (stars), and allY (triangles) calculated in simulations and estimated from experiments (by applying the law of rectilinear diameters and critical exponents^90^ to measurements of saturation concentrations at several temperatures^78^).The black solid line indicates perfect correlation between experimental and simulation results.

In contrast, the Urry hydrophobicity scale^88^ ranks the hydrophobicity of aromatic residues as W *>* Y *>* F. Consistently with this, the HPS-Urry model qualitatively captures the experimental trend in the relative phase separation propensities of the WT+NLS, allF, and allY variants, as indicated by the violet symbols in Fig.3. While the predicted order of phase separation propensities is correct, the absolute values of the critical solution temperatures predicted by the HPS-Urry model deviate significantly (by about *∼*20% on average) remaining consistently lower than the fitted experimental data r (Fig. 2C).

The HPS-cation–*π* model^56^ predicts higher-than-expected critical temperatures (approximately *∼* 30% higher on average) for the three variants (Fig. 2B and Fig. 3). This discrepancy can be attributed to a substantial overestimation of cation–*π* interactions in the HPS-cation–*π* model (Fig. A in SI). Importantly, the high discrepancy in the predicted critical temperature values result between the HPS-cation–*π* and HPS/HPS-Urry models highlights the prominence of cation–*π* interactions within A1-LCD condensates.

The CALVADOS2 model predicts coexistence lines that are, on average, very close to the experimental values (critical solution temperatures within approximately *∼* 5% of the values fitted by us from experiments). While the values are very close to the experimental ones, the relative ordering among the three variants deviates from the experimental trend (pink data in Fig. 3). It is important to note that the CALVADOS2 model was not specifically designed to predict critical solution temperatures, and this evaluation goes beyond its intended scope. In this context, a recent study by our group^89^, consistent with prior research^41,55,91^, emphasizes that models developed to capture single-molecule properties of IDRs do not necessarily reproduce condensate coexistence lines. While diverse parametrizations can yield reasonable predictions for radii of gyration for diverse IDRs, the accurate prediction of phase diagrams represents a far more rigorous and demanding validation test^89^.

The Mpipi model has demonstrated excellent performance in describing the phase behaviour of prion-like domain proteins^53,68^. Consistently, our tests show that Mpipi accurately predicts the order of the critical solution temperatures for the WT+NLS, allF, and allY variants (cyan symbols in Fig. 3). This agreement arises from the Mpipi model’s parametrization, which ranks the interactions of the three aromatic residues as W *>* Y *>* F. Despite its qualitative accuracy, Fig. 2E reveals that Mpipi slightly overestimates the critical solution temperatures for all three variants, with an average deviation of approximately 6%. Importantly, our analysis reveals that the improved Mpipi-Recharged model achieves the highest level of accuracy in reproducing experimental values, successfully capturing both the trend and the absolute critical solution temperatures for the three variants with deviations of less than *∼* 3% (green symbols in Fig. 3).

Overall, this analysis demonstrates that CALVADOS2, Mpipi, and Mpipi-Recharged offer a robust and accurate description of the condensate phase behaviour for A1-LCD aromatic mutants. In contrast, while the HPS-Urry model qualitatively captures the experimental phase-separation tendencies, it exhibits significant deviations in its quantitative predictions of critical solution temperatures.

### B. Comparison of the phase-separation saturation concentration between experiments and simulations

The concentration threshold above which biomolecular phase separation becomes thermodynamically favourable at a given temperature—C_*sat*_(T)—is one of the critical quantities that can be controlled by cells to trigger condensate formation and dissolution on demand^7,92^. In that sense, *in vitro* studies are extremely useful to characterise such threshold, characterising the border between a single homogeneous phase and the emergence of phase separated condensates^77,93–95^. Most of these studies have focused on measuring C_*sat*_ at physiological conditions of salt, pH, and temperature^22,24,93,95–100^. However, recent studies have also reported how C_*sat*_ varies with temperature and measured the highest temperature values beyond which phase separation is no longer achievable^77,78^. The saturation concentration quantifies the ability of a given protein to phase separate, thus, the higher the value, the lower propensity to form condensates^22,24,93,97^. Similarly, for condensates that present Upper Critical Solution Temperatures (herein critical temperature), the critical temperature evaluated through DC simulations serves as a direct indicator of the ability of a protein to undergo phase separation^69^. Indeed, the *in vitro* experimental values of saturation concentrations of the A1-LCD variants^77^ seem to correlate inversely with the critical solution temperatures that we can fit from the experimental coexistence densities^77^. A higher critical temperature is directly indicative of greater thermodynamic stability of the condensates. Thus, we can correlate the critical temperature obtained from simulations to the experimental saturation concentration at a given constant temperature (e.g., 298 K) to obtain a qualitative overview of the models sensitivity for describing protein phase behaviour as a function of specific sequence mutations^61,69^.

In Fig. 4, we show the simulated critical temperatures (normalised by the critical temperature of WT+NLS) against the experimental saturation concentrations^78^ measured at 298 K for the different A1-LCD mutants. The experimental C_*sat*_ vary significantly, changing by more than 2 orders of magnitude among the variants studied^78^. Thus, the comparison we perform in this section tests the sensitivity of the six different models to describe the impact of mutations on modulating the thermodynamic stability of A1-LCD condensates(Fig. 4A-F).

**FIG. 4.**
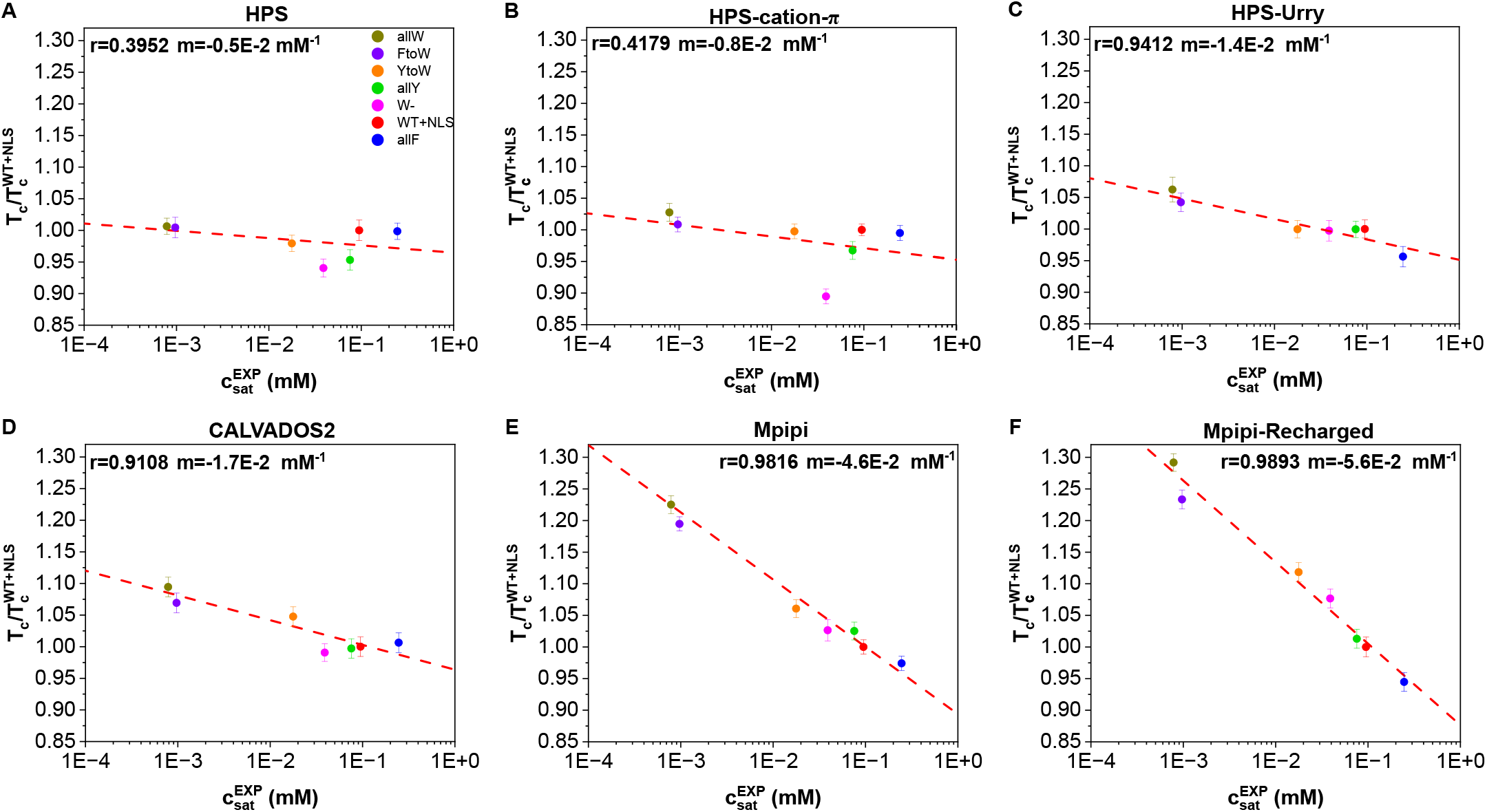
Predicted critical temperature (normalised by the critical temperature of the WT+NLS sequence) for the different models HPS (**A**), HPS-cation–*π* (**B**), HPS-Urry (**C**), CALVADOS2 (**D**), Mpipi (**E**), and Mpipi-Recharged (**F**) *vs*. the experimental saturation concentration at 298 K reported in Ref.^78^ for the different A1-LCD mutants. The panels include the Pearson correlation coefficient (*r*) and the slope (*m*) from a linear fit to the data. The error bars show the uncertainty in the critical temperature associated to its calculation using the laws of critical exponents and rectilinear diameters (see further details in Section V C).

To compare the predictions of the various models, we establish a linear relationship between the normalized critical solution temperatures and the experimental saturation concentrations, expressed as 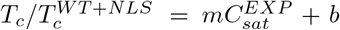, where *m* is the slope and *r* is the Pearson correlation coefficient that quantifies the quality of the fit. Notably, the HPS and HPS-cation–*π* exhibit the smallest absolute values for the slope *m* in this relationship, ranging from −0.5 × 10^*−*2^ to 0.8 × 10^*−*2^mM^*−*1^. This indicates that the critical temperatures predicted by these two models is only weakly affected by the mutations studied here (Fig. 4A-C, respectively). Furthermore, the quality of the fit is relatively low for the HPS (*r*=0.3952) and HPS-cation–*π* models (*r*=0.4179), suggesting a poor correlation between their predicted critical temperatures and the experimental trends for C_*sat*_.

The Mpipi and Mpipi-Recharged models predict the most significant changes in critical temperatures as a function of the variations in saturation concentration among the variants (Fig. 4E and 4F, respectively). This is reflected in the steep absolute values of their slopes, −4.6 × 10^−2^ and −5.6 × 10^−2^mM^−1^, respectively. Additionally, both models exhibit the strongest linear correlation with the experimental data, achieving Pearson correlation coefficients exceeding 0.98.

Finally, the HPS-Urry and CALVADOS2 models exhibit intermediate performance compared to the HPS/HPS-cation–*π* and Mpipi/Mpipi-Recharged models. The critical temperatures predicted by both HPS-Urry and CALVADOS2 display a strong linear correlation with the experimental C_*sat*_ values, achieving Pearson correlation coefficients above 0.91. However, both models predict only a moderate decrease in *T*_*c*_ as a function of changes in the saturation concentrations of the variants studied (Fig. 4C,D). Because the CALVADOS2 model was not specifically designed to predict critical solution temperatures, this evaluation extends beyond the model’s intended scope.

Building on these results, we now focus on the CALVADOS2, Mpipi, and Mpipi-Recharged models to further investigate the phase behaviour of A1-LCD mutants. Specifically, we use these models to compute the saturation concentration at 298 K (C_*sat*_) by extensively sampling the equilibrium protein concentration in the dilute phase in coexistence with the condensates through Direct Coexistence simulations. Estimating C_*sat*_ using this approach is computationally demanding due to the inherent challenges of sampling the dilute phase, particularly at low temperatures^68^. The dilute phase contains a very low concentration of molecules, meaning there are intrinsically few particles present. This low number of particles can result in significant variability in the measured concentrations across different regions of the simulation box, introducing substantial noise into the measurement. Furthermore, exchanges of molecules between the dilute and condensed phases are rare events, which require long simulation times to be captured adequately. These challenges are exacerbated at lower temperatures, where the equilibrium concentration in the dilute phase becomes even smaller. While computationally demanding, extracting C_*sat*_ from the DC method allows us to perform a direct comparison between the same observable measured in the simulations and the experiments.

In Fig. 5, we present the predicted saturation concentration values from simulations, normalised by the saturation concentration of the WT+NLS sequence, plotted on a logarithmic scale as is customary for saturation concentrations, 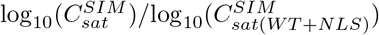. The predictions from the CALVADOS2, Mpipi, and Mpipi-Recharged models are compared against experimentally determined normalised saturation concentrations for the studied mutants 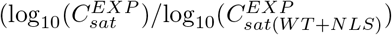 Section V F for further information) reported in Ref.^78^. To quantify the agreement between the simulations and experiments, we calculate *D*, a metric that measures the average deviation of the simulation predictions from the experimental values (details provided in Methods). Lower *D* values indicate closer agreement with the experimental data. Additionally, we perform a linear fit to examine the relationship between the normalised simulation results and the experimental data (red dashed line in Fig.5), comparing it to the perfect fit (diagonal black line in Fig.5). To assess the strength of the linear correlation, we calculate the Pearson correlation coefficient (*r*). This metric evaluates whether the models qualitatively capture the experimental trends, even if quantitative deviations are present.

**FIG. 5.**
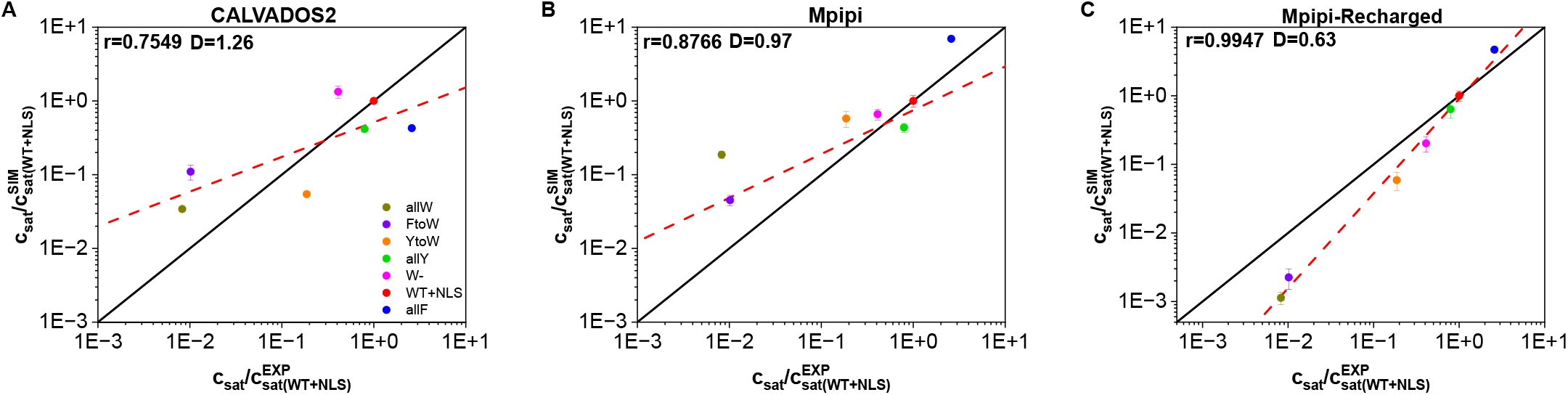
Simulated *vs*. experimental saturation concentration (C_*sat*_), normalised by the WT+NLS C_*sat*_^78^ for the different variants and for the models CALVADOS2 (**A**), Mpipi (**B**) and Mpipi-Recharged (**C**). The Pearson correlation coefficient (*r*) and the root mean square deviation from the experimental values (*D*) are displayed for each set of modelling data. The black lines indicate a perfect match between experimental and computational values, while the red dashed lines depict a linear regression for each set of data.

We find that the prediction of Mpipi-Recharged model for the normalized critical solution temperature presents the lowest average error, *D* = 0.63, with respect to the experimental values and the highest Pearson correlation coefficient among the set (*r* = 0.9947), indicating exceptional agreement with the experimental trends. The Mpipi and CALVADOS2 models also provide good de-scriptions with mean errors of *D* = 0.97 and *D* = 1.26, respectively, and Pearson correlation coefficients of *r* = 0.8766 and *r* = 0.7549, respectively.

In addition, we compare the absolute experimental and simulation log10(*C*_*sat*_) values (Fig. B in SI). In this comparison, the values of the Person correlation coefficients remain unchanged from those in Fig.5. That is, the predictions of the Mpipi-Recharged model exhibit the strongest correlation with the experimental values of log10(*C*_*sat*_), followed by Mpipi, and finally CALVADOS2. In contrast, he size of the mean errors, *D*, do change, revealing that CALVADOS2 provides log10(*C*_*sat*_) with the smallest overall errors (*D* = 0.38), followed by Mpipi-Recharged (*D* = 0.78) and finally Mpipi (*D* = 0.93).

Overall, our comprehensive evaluation in this Sectionhows that all three models—CALVADOS2, Mpipi, and Mpipi-Recharged—perform exceptionally well in predicting the phase-separation propensity of A1-LCD variants, with Mpipi-Recharged standing out as the most accurate. Their excellent performance is particularly noteworthy given the substantial challenges coarse-grained models face in reproducing experimental coexistence lines.

### C. Condensate viscoelastic behaviour of A1-LCD mutants by different models

Condensate viscoelastic properties have been linked to the roles of these systems in health and disease^34,101,102^. Whereas liquid-like states in condensates are associated to biological function, their progressive transition into solid and gel-like states has been linked to the onset of multiple neurological disorders^24^. Examples of protein condensates displaying hardening over time have been reported for RNA-binding proteins such as FUS^103,104^, TDP-43^105^, or hnRNPA1^106^ among others^30^. Hence, it is important to determine the ability of different residue-resolution CG models to probe the material properties of condensates: i.e., whether they behave as liquids or gels, and whether their viscosity remains constant or increases over time. Moreover, determining the ability of residue-resolution CG models to characterise which precise interactions and domains control their time-dependent transport properties^103,107–109^ is central. Such information enables to understand the molecular onset between functional and dysfunctional behaviour.

Computationally, the generalised Green-Kubo relation^110,111^ (see Section V E) is one of the most efficient methods for evaluating viscosity of protein condensates using residue-resolution CG models^31,69,112^. Calculating the shear stress relaxation function (*G*(*t*)) using the Green-Kubo approach provides information of both inter- and intra-molecular protein interactions, which may vary significantly with the chosen CG model^69,108,112^. Since the predictive accuracy of most residue-resolution CG models has been optimized and tested using experimental single-protein radius of gyration and phase diagrams (either saturation concentrations or critical solution temperatures)^41,43,53,54,59^, evaluating their ability to predict condensate viscosity provides a demanding benchmark that extends beyond the scope of their original parameterisation.

As the experimental reference, we use the experimental *in vitro* viscosity measurements at *T ∼* 298 K from Ref.^78^ for the seven variants of A1-LCD studied in the previous sections. We only test the performance of the CALVADOS2, Mpipi, and Mpipi-Recharged models because 1) HPS and HPS-cation–*π* failed to recapitulate the relative phase-separation propensity of the A1-LCD variants (Fig. 4A-B, respectively); and 2) HPS and HPS-Urry do not phase-separate at 298 K, where condensates experimental viscosities have been reported.

To estimate the viscosity, we calculate the shear stress relaxation function, *G*(*t*), of the different A1-LCD variants under bulk conditions at 298 K (see Fig. 6A and Supporting Movie B for representative configurations of such simulations, and Section V B for the simulation details). For these calculations, we perform bulk simulations (in the canonical ensemble) of ∼ 3 – 5*μ*s depending on each specific system as reported in Section V B.

**FIG. 6.**
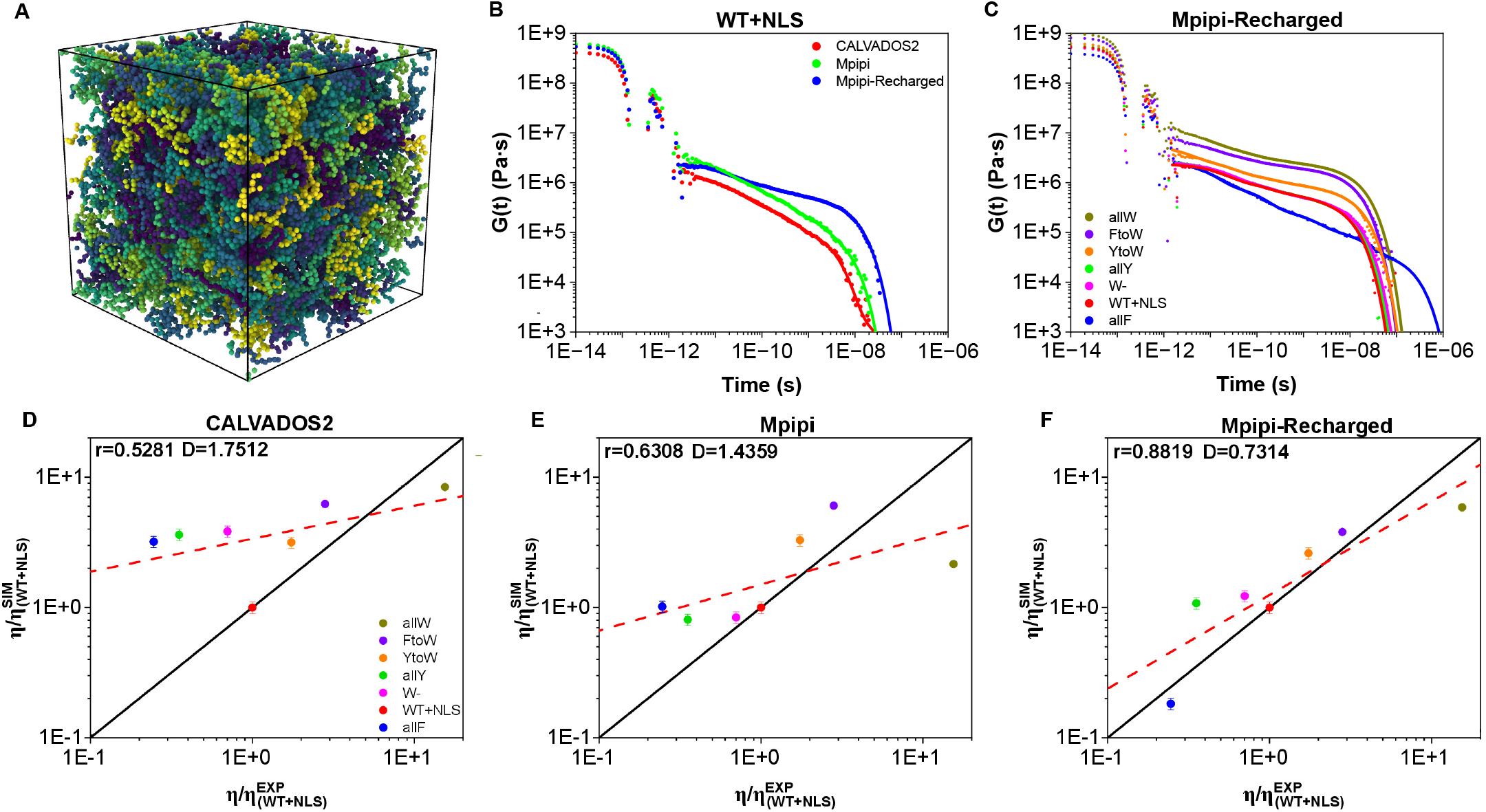
(**A**) Snapshot of a bulk NVT simulation with 200 A1-LCD (WT+NLS) protein replicas (each of them depicted by a different tone of colour) employed to compute the condensate viscosity. (**B**) Shear stress relaxation modulus (*G*(*t*)) for the WT+NLS sequence at 298 K evaluated by different models. (**C**) *G*(*t*) for different A1-LCD mutants using the Mpipi-Recharged model. In both **B** and **C** panels, *G*(*t*) raw data obtained during the simulations are represented by filled symbols, while Maxwell’s mode fits to the data (as described in Section V E) are plotted with solid lines. **D, E** and **F**: predicted versus *in vitro* viscosity (both reduced by the corresponding WT+NLS value) for all variants under study and the CALVADOS, Mpipi and Mpipi-Recharged models respectively. The meaning of *r, D*, and the red dashed and black solid lines is the same as in Fig. 5.

The calculation of *G*(*t*) for the WT+NLS sequence using the three models is shown in Fig. 6B. While the predictions of the three models for *G*(*t*) at short timescales are rather similar, indicating that bond, angle, and intramolecular conformational relaxation is similar for all of them, we observe large differences at long times. At long timescales, the slope and terminal decay of *G*(*t*), which strongly determines the condensate viscosity, differs significantly among the three models, clearly demonstrating the critical influence of the model parameterisation on the resulting viscoelastic properties of the condensates. Such variability is even more pronounced when comparing the behaviours of *G*(*t*) predicted by the three models for all the different A1-LCD mutants. We show the full *G*(*t*) plots predicted by the Mpipi-Recharged model in Fig. 6C, and the viscosity values for the three models in Fig. 6D-F. The calculation of *η* from *G*(*t*) consists in integrating the shear stress relaxation modulus (*G*) as a function of time (for further details see Section V E).

Fig. 6C reveals that the amino acid mutations tested lead to drastic variations in the condensate viscosity predicted by Mpipi-Recharged (of approximately two orders of magnitude), in perfect agreement with the experimental observations^78^. This results highlights the strong influence that specific sequence mutations have on regulating the transport properties of condensates, as previously found for FUS^113^, TDP-43^105^, and Tau^114,115^.

We next compare the viscosity *η* values for all A1-LCD sequences as predicted by simulations using the CALVA-DOS2 (Fig. 6D), Mpipi (Fig. 6E), and Mpipi-Recharged (Fig. 6F) models against the corresponding experimental values^78^. Both simulation and experimental viscosity values are normalised by the WT+NLS *η* value to enable a relative comparison. This normalisation is necessary because the absolute values predicted by all the residue-resolution coarse-grained models systematically underestimate the experimental viscosities by several orders of magnitude^69^. This discrepancy arises naturally from the implicit treatment of solvents and ions, as well as the neglect of atomic vibrations and detailed inter-atomic interactions among amino acids, due to the amino acids being represented as spherical beads. While the choice of the A1-LCD WT+NLS variant used to normalise the viscosities influences the absolute deviation between simulations and experiments, it does not alter the overall trend.

We observe that the sensitivity exhibited by the different models in their predictions of critical temperatures (Fig.4D-F) is similarly reflected in the variations in condensate viscosity (Fig.6D-F). The Mpipi-Recharged model exhibits the greatest sensitivity, with viscosity values spanning nearly two orders of magnitude among the different A1-LCD variants, and emerges as the best fit for capturing the viscoelastic behaviour of A1-LCD condensates. Specifically, the Pearson coefficient (*r*) assessing the strength of the correlation between simulation and experimental *η* values is high for Mpipi-Recharged (*r* = 0.8819), and moderate for Mpipi(*r* = 0.6308) and CALVADOS2 (*r* = 0.5281). In addition, the root mean squared deviation (*D*) describing the mean error from the experimental data is smallest for Mpipi-Recharged (*D* = 0.7314), and moderate for Mpipi (*D* = 1.4359) and CALVADOS2 (*D* = 1.7512).

We also investigate whether a correlation between the critical temperature and condensate viscosity can be established for the different studied variants. A correlation between these two quantities is expected since intermolecular interaction strength, which favours condensation (i.e., increases T_*c*_), should also diminish molecular mobility (i.e., increasing *η*). In Fig. 7, we plot the critical temperature (normalised by the T_*c*_ of the WT+NLS sequence) *vs*. condensate viscosity (also normalised by the *η* of the WT+NLS sequence) for the different studied A1-LCD mutants as predicted by CALVADOS2 (A), Mpipi (B), and Mpipi-Recharged (C) models. Moreover, experimental *in vitro* data from Ref.^78^ for the WT+NLS, allF and allY sequences are included (empty squares). As it can be seen, the Mpipi-Recharged predicts the correlation experimentally found for these three variants (depicted by a grey band linearly extrapolated). The full phase diagram (including T_*c*_) for the rest of the variants was not reported in Ref.^78^, nevertheless, it is expected that a similar correlation as that predicted by the Mpipi-Recharged model holds since it reasonably predicts the experimental critical temperature, saturation concentration, and condensate viscosity of all these variants, as shown in Figs. 2, 5, and 6, respectively. The CALVADOS2 also shows a correlation between the critical temperature and viscosity (Fig. 7A). However, the CALVADOS2 predictions do not span the same range of values for viscosity as the experimental measurements (Fig. 6D). For the case of Mpipi, a weak correlation between these the critical solution temperature and the viscosity is observed (Fig. 7B).

**FIG. 7.**
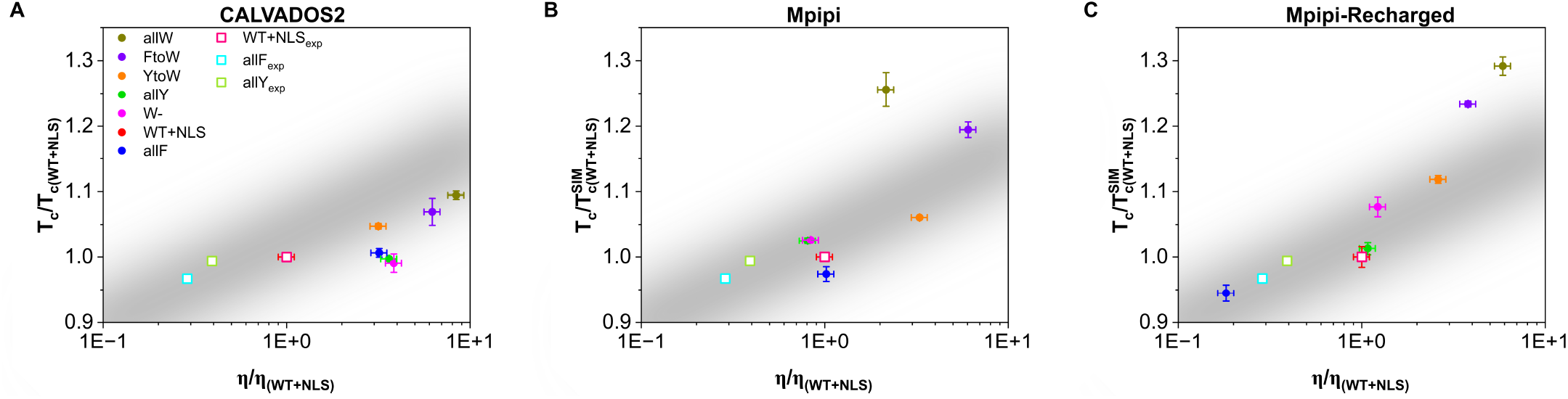
Correlation between the critical temperature (normalised by the T_*c*_ of the WT+NLS sequence) and the condensate viscosity (also normalised by the *η* of WT+NLS) for the different studied A1-LCD mutants as predicted by the CALVADOS2 (**A**), Mpipi (**B**), and Mpipi-Recharged (**C**) models. Experimental data from Ref.^78^ for the WT+NLS, allF and allY sequences have been also included as empty squares (see legend). A linear trend to the experimental correlation between these two quantities for the three sequences measured *in vitro*^78^ is shown with a grey band.

Future work exploring whether a correlation between the critical solution temperature and the viscosity arises in other condensates would be highly valuable, as both properties are challenging to measure and are rarely reported under identical conditions^77,78^. Establishing such a relationship could enable the qualitative inference of one property from the other, offering complementary insights into the stability and transport properties of biomolecular condensates.

### D. Key intermolecular interactions in A1-LCD condensates

Computational approaches can contribute to a deeper understanding of biomolecular phase separation by evaluating the underlying molecular information that is hardly accessible in experiments^69,74,116^. A powerful example is the calculation of intra and inter-molecular contact maps^52,69,89^ which provide microscopic insights about the key amino acids and protein domains enabling biomolecular self-assembly. However, the outcome of this type of analysis is intimately related to the specific model characteristics and its interaction parameters. Therefore, different models may provide widely different contact domain probabilities for the same protein condensate.

Analysing the differences in the inter-molecular contact maps predicted by the different the models we benchmark here, also allows us to understand the reasons why such models present different performance in these benchmarks. Specifically, we count the number of times a pair of amino acids belonging to different proteins within the condensate are in contact, where ‘in contact’ is defined as being closer than a cutoff distance which depends on the identity of the amino acids in the pair (see Section V D). We then evaluate the energy contribution of each contact considering the sum of the potential energy terms in the model at the interacting distance. That is, the sum of the Wang-Frenkel and Yukawa potentials for the Mpipi-Recharged model, Wang-Frenkel and Debye-Hückel for Mpipi, and Lennard-Jones and Debye-Hückel for CALVADOS2 and the HPS family (see Section V D). The resulting contact energy maps reveal the key molecular interactions that sustain the liquid network within the different condensates.

When comparing the contact energy maps for WT+NLS condensates predicted by the different models at 0.95 T_*c*_ (Fig. C in SI), we find significant differences among the behaviours of the various models. The HPS and HPS-Urry models present homogeneous maps of intermolecular interactions, with most amino acid pairs presenting contact energies that differ by at most 0.002 kJ/mol. In contrast, the HPS-cation–*π*, Mpipi and Mpipi-Recharged models give rise to heterogeneous maps, where the contact energies among amino acid pairs can differ by more than 0.004 kJ/mol. CALVA-DOS2 presents an intermediate behaviour between the two extremes. Such difference can be explained by the HPS and HPS-Urry models considering electrostatic and *π*-*π* interactions as stronger contributors to biomolecular phase separation than cation–*π* contacts (see Fig. D in SI, and the relative interaction strength maps in Fig. A in SI). In the CALVADOS2 model, such a difference is present, but is much more moderate^55^. The HPS-cation–*π*, Mpipi, and Mpipi-Recharged models predict that phase separation is most strongly contributed by some specific prion-like sub-domains. Whilst all models show a strong interacting region from the 105th to 115th residue (a sub-domain enriched in Y, F, and G), the HPS-cation–*π*, Mpipi, and Mpipi-Recharged also present a highly interacting sub-domain from the 130th amino acid to the 135th residue, which contains 2 arginines and 1 phenylalanine. In addition, the HPS and HPS-cation–*π*, but more prominently the Mpipi, and Mpipi-Recharged models show two sub-domains between the 75th and 85th position (enriched in asparagine and aromatic residues) and between the 15th and 45th (rich in N, F, R and G) that significantly contribute to the stability of A1-LCD condensates through cation-*π* and *π*-*π* interactions (Fig. C in SI).

Since the Mpipi-Recharged model has consistently emerged as the most accurate model of those we have tested (Figs. 1-6), we now employ it to compare the impact of amino acid mutations on the contact energy maps of A1-LCD condensates. In Fig. 8, we present the contact energy maps for condensates of the WT+NLS, allF, and allW sequences at 0.95 T_*c*_, where T_*c*_ represents the critical temperature specific to each variant (Fig.8A-C). Notably, the contact energy maps for the allF and WT+NLS condensates exhibit striking similarities (Fig.8A and 8B). In contrast, the contact energy map for the allW condensates (Fig.8C) reveals significantly stronger contact interactions among residues near the protein terminal region, particularly from the 90th residue onward. This increase in intermolecular contacts at the protein terminal region enhances the interaction energies across the whole protein sequence, suggesting a cooperative mechanism that reinforces the condensate assembly (Fig. 8C). This increase of intermolecular interactions in the allW mutant explains the large increase in viscosity observed both computationally and experimentally (Fig. 6) for this variant with respect to the WT+NLS and allF sequences.

**FIG. 8.**
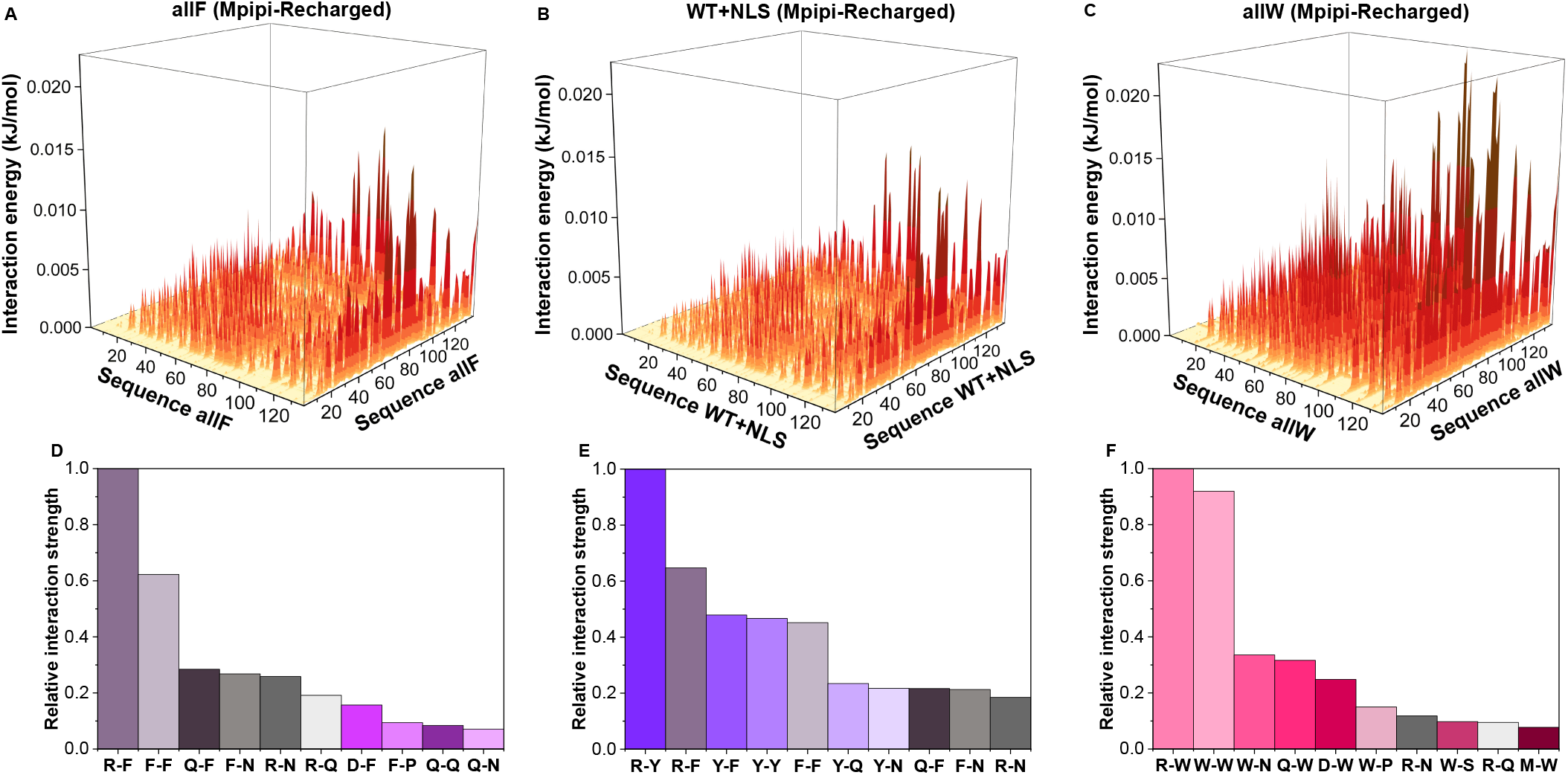
Contact energy maps of pairwise intermolecular interaction energy for allF (**A**), WT+NLS (**B**), and allW (**C**) condensates at T = 0.95 T_*c*_ as predicted by the Mpipi-Recharged model. The most frequent intermolecular amino acid pairwise interactions sustaining the condensates of allF (**D**), WT+NLS (**E**), and allW (**F**) are also displayed. Pairwise contacts have been both normalised by the highest value of residue-residue pairwise interactions and their abundance across their sequence (see Section V D for further details on these calculations).

We now extract from Fig. 8A-C the most frequent amino acid pairs contributing that stabilise the condensed phase, as predicted by the Mpipi-Recharged model (Fig. 8D-F). The relative interaction strength of each pair has been normalised by that of the contact pair with highest energetic contribution. The relative interaction strength of each pair has been normalized to the contact pair with the highest energetic contribution. As expected, the key intermolecular interactions in WT+NLS condensates are primarily cation–*π* (e.g., R–Y and R– F) and *π*–*π* interactions (e.g., F–F, Y–F, and Y–Y). In allF condensates, the dominant contacts include cation-*π* (R–F) and *π*–*π* interactions (Y–F and F–F), along with interactions involving asparagine (N) with arginine (R) and phenylalanine (F). For allW condensates, the most prevalent interactions are W–W and W–R, followed by contacts between tryptophan (W) and asparagine (N), glutamine (Q), and aspartic acid (D).

These findings align with the stickers-and-spacers framework for protein phase transitions, in which multivalent proteins are conceptualized as heteropolymers comprising ‘stickers’ (binding sites for associative interactions) and ‘spacers’ (regions between stickers)^117,118^. Within the stickers-and-spacers framework, as proposed in Ref.^77^ for A1-LCD condensates, tyrosine (Y), phenylalanine (F), and tryptophan (W) function as the primary stickers, arginine (R) acts as a context-dependent sticker, and other amino acids serve as spacers. Our simulations demonstrate that the predictions of the Mpipi-Recharged model are in excellent agreement with this framework (Fig. 8D-F).

We now focus on the most frequent interactions in WT+NLS condensates, as predicted by the Mpipi-Recharged (Fig. 8E) and the CALVADOS2 models (Fig. E2D). While the Mpipi-Recharged model suggests that cation–*π* interactions such as R–Y and R–F are stronger contributors than Y–Y, F–F, and Y–F to the stability of A1-LCD condensates, CALVADOS2 proposes that instead *π*–*π* contacts are more energetically favourable than cation–*π* interactions for these systems (Fig. DD in SI). As in the Mpipi-Recharged model, according to the Mpipi model, the five most energetically favourable contacts in A1-LCD condensates are R–Y, R–F, Y–F, F–F, and Y–Y (Fig. DE in SI). In contrast, the HPS and HPS-Urry models do not predict cation–*π* interactions as main contributors for A1-LCD phase-separation (Fig. DA,C in SI). Instead, HPS and HPS-Urry consider that electrostatic R–D and K–D contacts alongside *π*–*π* interactions (such as Y-F, F-F, and Y-Y, which in the model parameterisation have a considerable interaction strength^76^, see Fig. A in SI) are the key contacts driving biomolecular phase separation. The underestimation of cation–*π* interactions for biomolecular phase transitions was addressed by the HPS-cation–*π* reparametrisation^56^, which added an extra potential term to the model (see Section V A for further details). Nevertheless, as shown in Fig. DB in SI, the HPS-cation–*π* reparametrisation significantly overestimates cation–*π* contacts predicting that R–Y, R–F, K–Y and K–F contribute four times more than aromatic interactions such as F–F or Y–F. Such overestimation of cation–*π* interactions by the HPS-cation–*π* model leads to very high critical temperatures for A1-LCD condensates, as shown in Fig. 2B; this has been previously discussed in Refs.^31,53,56,69,87^ for other RNA-binding proteins such as FUS, TDP-43, or DDX4, among others.

Overall, this analysis reveals that residue-resolution CG models considering both cation–*π* and *π*–*π* interactions as primary contributors to the stability of A1-LCD condensates align most closely with experimental observations. Notably, models such as Mpipi and Mpipi-Recharged, which emphasize the stronger contribution of cation–*π* interactions over *π*–*π* interactions, show the highest level of agreement with experimental data.

## IV. DISCUSSION

Computer simulations are a powerful tool to complement the *in vitro* and *in vivo* investigations of biomolecular condensates^42,119^. In this study we have compared the capacity of six different sequence-dependent residue-resolution CG models to predict temperature-vs-concentration phase diagrams, critical solution temperatures, saturation concentrations, and viscosities of A1-LCD biomolecular condensates. We systematically compare the predictions of the various models against experimental coexistence lines, critical solution temperatures fitted from experimental coexistence densities, experimentally measured saturation concentrations, and experimentally measured viscosities for A1-LCD condensates, and seven mutants. Our results show that the Mpipi-Recharged model displays quantitative accuracy in predicting the critical temperature and condensate saturation concentration of the different A1-LCD mutants (Figs. 2 and 5) reported in Ref.^77,78^.

Furthermore, evaluating condensate viscosity provides a stringent test of the ability of these models to predict the biophysical properties of condensates beyond stability. Our analysis reveals that the CALVADOS2, Mpipi, and especially the Mpipi-Recharged models accurately capture the relative variations of the viscoelasticity of protein condensates as a function of specific sequence mutations (Fig. 6). This highlights the potential of residue-resolution coarse-grained models to effectively predict how sequence variations influence the material properties of condensates beyond the systems tested here.

Importantly, characterizing viscoelasticity is essential for distinguishing between liquid-like and solid-like behaviours of condensates, which are closely tied to biological functions and pathological malfunctions^24,34,42,101,102^. Notably, we observe a clear correlation—both emerging from the experimental and computational data (Fig. 7)—between condensate thermodynamic stability (i.e., T_*c*_) and viscoelasticity. This correlation suggests that higher critical temperatures for phase separation are linked to increased condensate viscosities. This relationship emerges because both greater stability and viscosity are promoted by stronger intermolecular interactions: stronger biomolecular interactions promote condensation and, simultaneously, reduce molecular diffusivity, thereby enhancing viscosity^31,103^.

Finally, we have evaluated the intermolecular contact energy maps of A1-LCD condensates as predicted by the different models studied here, paying particular attention to the Mpipi-Recharged force field (Fig. 8). Our analysis explain how the variations in the predictions of the different models may be explained by the differences in the the pairwise residue-residue interactions that sustain A1-LCD condensates.

The best-performing models in our benchmarks— namely CALVADOS2, Mpipi, and the Mpipi-Recharged model—consider in their parametrizations that both cation–*π* interactions (e.g., R–Y and R–F) and *π*–*π* contacts (e.g., F–F, Y–F, and Y–Y) are the dominant intermolecular forces stabilising A1-LCD condensates. Among these models, Mpipi-Recharged, followed by Mpipi, stands out as the most accurate, particularly because they rank cation–*π* interactions as stronger contributors than *π*–*π* contacts. In contrast, models that underestimate the contribution of cation–*π* interactions—such as the HPS and HPS-Urry models—perform less accurately, failing to capture the significant shifts in critical temperatures and saturation concentrations caused by aromatic residue mutations (Fig. D in SI). While the HPS-cation–*π* model overhauls the contribution of cation–*π* interactions, it overemphasizes their strength, resulting in predictions of unrealistically high critical temperatures (Fig.2).

A closer examination of the predictions made by the Mpipi-Recharged model reveals that, in addition to capturing the importance of cation–*π* and *π*–*π* contacts, it also identifies prevalent interactions involving aromatic residues (Y and F) and arginine with asparagine (N) and glutamine (Q) within A1-LCD condensates. These predictions are consistent with experimental phase diagrams of A1-LCD systems and align with the stickers-and-spacers framework, which demonstrates that tyrosine, phenylalanine, and tryptophan act as primary stickers, arginine serves as a context-dependent sticker, and other amino acids function as spacers^77,117^.

Recent advancements in residue-resolution coarsegrained models for biomolecular phase separation have led to significant progress. Beyond the examples presented here, models such as the HPS-Urry model, the CALVADOS family, and the Mpipi family have demonstrated exceptional predictive power in describing the relative thermodynamic stability of a wide range of biomolecular condensates, as well as single-molecule observables of proteins, in agreement with experimental data. Although the HPS model does not perform particularly well in our benchmarks, it is important to highlight that all other models were built upon the foundational framework provided by the pioneering HPS model^41,52,56^. Thus, it is unsurprising that these newer models outperform the original HPS model, reflecting the continuous progress in the field.

There are now many more sequence-dependent residue-resolution CG models, such as the original CALVA-DOS^41^, KH^86^ or the FB-HPS^58^, which were not included in this work due to computational constraints, which limit the number of models feasible to simulate. However, as demonstrated in our previous study^53^, these models perform reasonably well in capturing the phase diagrams of A1-LCD condensates and their mutants.

With this study, we aim to highlight the continuous advancements in residue-resolution coarse-grained model development, which is increasingly enabling the quantitative description of biomolecular phase behaviour while providing complementary microscopic insights that are often challenging to obtain through traditional experimental methods.

## V. METHODS

### A. Model and methods

In our simulations, we represent the intrinsically disordered regions (IDRs) as coarse-grained structures with one bead per amino acid. The potential energy of the coarse-grained force field is defined as follows:

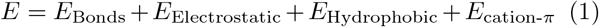

**TABLE S1.** Table of equations used by each model. The details of the equations are given below.

| Model \ Energy | $E_{\text{Bonds}}$ | $E_{\text{Electrostatic}}$ | $E_{\text{Hydrophobic}}$ | $E_{\text{cation-}\pi}$ |
| --- | --- | --- | --- | --- |
| HPS | (2) | (3) | (7) | - |
| HPS-cation-π | (2) | (3) | (7) | (10) |
| HPS-Urry | (2) | (3) | (7) | - |
| CALVADOS2 | (2) | (3) | (7) | - |
| Mpipi | (2) | (3) | (8) | - |
| Mpipi-Recharged | (2) | (6) | (8) | - |

Bonded interactions are modeled by a harmonic potential

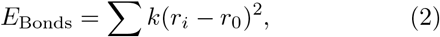

where *k* = 9.6kJ/mol *·* Å^2^ for all the models except for the HPS-cation-*π* which is *k* = 2.4kJ/mol *·* Å^2^ and for the HPS-Urry which is *k* = 4.8kJ/mol *·* Å^2^. The equilibrium bond length is *r*_0_ = 3.81 Å between bonded amino acid beads.

The electrostatic interactions (*E*_Electrostatic_) among charged amino acids are governed by a Debye-Hückel potential

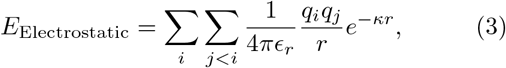

where *q*_*i*_ and *q*_*j*_ represent the charges of the beads *i* and *j, ϵ*_*r*_= 80 *ϵ*_0_, is the relative dielectric constant of water (being *ϵ*_0_ the electric constant), *r* is the distance between the *i*th and *j*th beads, and *κ*= 1 nm^−1^ is the Debye screening length that mimics the implicit solvent (water and ions) at physiological salt concentration (*c*_*s*_ ∼ 150mM of NaCl for the models HPS, HPS-cation-*π*, and the HPS-Urry model, and for the Mpipi model the value is *κ*= 1.26 nm^−1^. For the CALVADOS2 and Mpipi-Recharged model *κ* depends on the salt concentration as

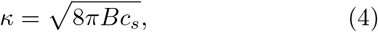

where *B*(*ϵ*_*r*_) = *e*^2^*/*4*πk*_*B*_*Tϵ*_0_*ϵ*_*r*_ is the Bjerrum length. The relative dielectric permittivity depends on the temperature according to this empirical law

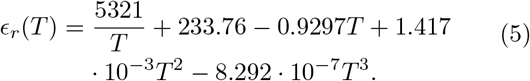

For the Mpipi-Recharged model, electrostatic interactions are modelled through a Yukawa potential given by

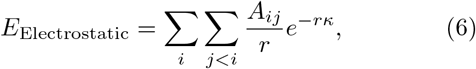

where *A*_*ij*_ controls the interaction between a pair of amino acids and *κ* modulates the salt concentration in an explicit way. The dielectric constant varies with temperature according to Eqs. (4) and (5).

For the HPS, HPS-cation-*π*, HPS-Urry and CALVA-DOS2 models, the hydrophobic interactions are established using a scale of hydrophobicity derived from statistical analysis of amino acid contacts in PDB structures. These interactions are incorporated using the Ashbaugh-Hatch potential^43,86,120,121^ functional form given by

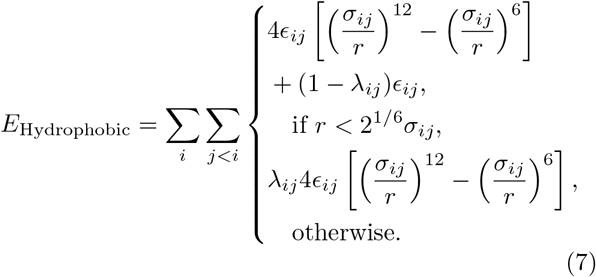

where *λ*_*i*_ and *λ*_*j*_ are parameters that account for the hydrophobicity of the *i*th and *j*th interacting particles respectively, being *λ*_*ij*_ = (*λ*_*i*_ + *λ*_*j*_)*/*2 following the Lorentz-Berthelot mixing rules^122,123^. The excluded volume of the different residues is given by *σ*_*i*_ and *σ*_*j*_, where *σ*_*ij*_ = (*σ*_*i*_ +*σ*_*j*_)*/*2, and *r* is the distance between the *ij* particles. The parameter *ϵ*_*ij*_ is set to 0.2 kcal/mol to reproduce experimental single-IDR radius of gyration^43^.

In the Mpipi and Mpipi-Recharged models, the hydrophobic interactions are parameterized by the Wang-Frenkel potential given by

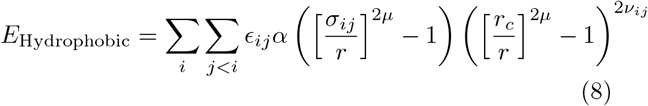

where

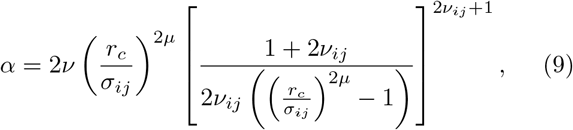

where the excluded volume of the different residues is given by *σ*_*ij*_, *r* is the distance between the *ij* particles, *ϵ*_*ij*_ is the energy interaction parameter, *r*_*c*_ = 3*σ*_*ij*_ is the cut-off of the potential between those amino acids and *µ*=1 and *ν*_*ij*_ are terms that are involved in the shape of the potential. The values of *σ*_*ij*_ and *ϵ*_*ij*_ are precisely parametrize for each interaction^61,79^

For the HPS-cation-*π* model another additional component is added. A reparametrisation of the interactions between the positively charged amino acids and the aromatic ones as a Lennard-Jones potential is modelled as

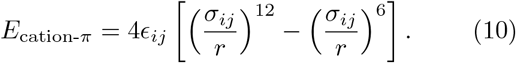

We employed a cut-off of 3*σ*_*ij*_ for the hydrophobic interactions, and 3.5 nm for the electrostatic ones^43^, except for the HPS-Urry that uses a cut-off of 2.0 nm and for the CALVADOS2 that uses a cut-off of 4.0 nm^55^.

In table S1 we provide a summary of the different potentials used by each model studied in this work. To better visualize the differences in interactions between the different models, Fig. A in SI shows the normalized values of interaction strength for all amino acid pairs. Higher values indicate stronger attractive interactions, and lower values indicate weaker or repulsive interactions.

### B. Preparation of the systems and simulation details

All simulations were conducted using the LAMMPS software^124,125^. Both Direct coexistence (DC) simulations and viscosity calculations were performed in the NVT ensemble employing a Langevin thermostat^126^ with a relaxation time of 5 ps and a time step of 10 fs for the Verlet integration. The configurations for DC simulations were generated by placing 200 protein replicas in a slab with ∼17×17nm^2^ of section and 120nm of length for a resulting average density of ∼0.1 g/cm^3^. Simulations to calculate the coexistence densities run a total time of the order of ∼1*µ*s after reaching equilibrirum. Calculation of the viscosity was carried out from NVT simulations by placing 200 protein replicas in a cubic box that was isotropically compressed to reach the desired bulk density. After equilibration, production run took around ∼3-5*µ*s to fully capture the stress relaxation of each specific system.

### C. Direct Coexistence technique

Employing Direct Coexistence (DC) simulations^84,127–129^, we determined the phase diagram for each of the studied proteins^78^. This method involves simulating the two coexisting phases within the same simulation box. In our approach, we arranged a high-density protein liquid alongside a very low-density counterpart. To accommodate the diverse densities, we employed an elongated simulation box, therefore allowing for the formation of both coexisting phases forming an interface perpendicular to the long direction of the simulation box. Equilibrium was attained through NVT simulations, and subsequently, we measured the equilibrium coexisting densities of both phases along the elongated side of the box. This process was repeated across various temperatures until reaching the critical temperature. To mitigate finite system-size effects near the critical point, we calculated the critical temperature (*T*_c_) and density (*ρ*_c_) using the law of critical exponents and rectilinear diameters^130^

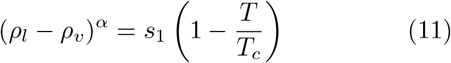

and

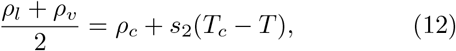

where the notation *ρ*_*l*_ and *ρ*_*v*_ are the densities of the condensed and diluted phases, respectively. Moreover, s_1_ and s_2_ are fitting parameters, while the critical exponent, *α* = 3.06 for the three-dimensional Ising model^130^. The critical temperature error has been determined using the law of critical exponents and rectilinear diameters to estimate the system’s critical temperature by considering the last two, three, and four temperatures.

### D. Calculating contact maps and top contacts

The computation of intermolecular and intramolecular contact maps within protein condensates is calculated from DC trajectories. Contacts were determined across all systems at temperatures approximately 0.95*T*_c_, where *T*_*c*_ corresponds to the critical temperature of each system. Typically, molecular contacts are identified based on a distance criterion, with the assumption that the relative frequency of contact map occurrences (rather than absolute frequency) remains generally unaffected by the selected cut-off distance used in calculations, provided the cut-off values are reasonable. However, to accurately capture the most relevant and common residue-residue contact pairs that facilitate LLPS, it is highly recommended to account for the specific parameterisation of each amino acid in terms of excluded volume and minimum potential energy interaction distance. Hence, we adopted a sequence-dependent cut-off distance equivalent to 1.2*σ*_*ij*_, where *σ*_*ij*_ represents the mean excluded volume of the respective ith and jth amino acids^69^. Given that the minimum of the potential used lies at approximately 2^1*/*6^*σ*_*ij*_ ≈ 1.122*σ*_*ij*_, we set the cut-off distance slightly beyond this point, at 1.2*σ*_*ij*_, to ensure significant binding. By implementing this innovative sequence-dependent cut-off scheme for each amino acid pair interaction, we can effectively filter out adjacent contacts that may coincide with actual interacting amino acids along the sequence, thus enhancing our ability to accurately identify the amino acids that positively contribute to stabilising condensates^69^.

To define the relative interaction strength for the most relevant contacts, this interaction frequency described above (*f*_i,j_) is weighted with the hydrophobic and electrostatic contributions of each amino acid pair for the model 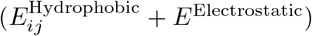, normalized against the occurrence frequency of each pair of amino acids in the sequence, as described by

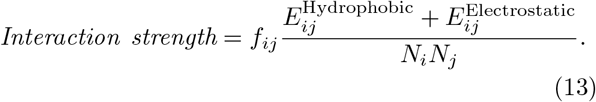

### E. Calculating viscosities

We can determine the viscosity of the LCD model’s condensate using the Green-Kubo (GK) relation. The time-dependent mechanical response of a viscoelastic material under small shear deformation is characterized by the shear stress relaxation modulus (*G*(*t*))^131^. When subjected to zero deformation, *G*(*t*) can be computed by correlating any off-diagonal component of the pressure tensor at equilibrium^132,133^

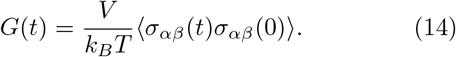

In Eq.14, *σ*_*α*,*β*_ represents an off-diagonal component of the stress tensor, *V* is the volume, *T* is the system temperature and *k*_*B*_ is the Bolztmann’s constant. For isotropic systems, a more precise expression of *G*(*t*) is obtained by considering the six independent components of the pressure tensor^112,134–136^, given by

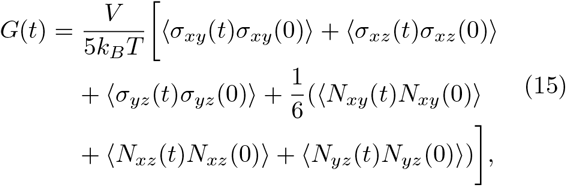

where *N*_*α*,*β*_ = *σ*_*α*,*α*_ − *σ*_*β*,*β*_ represents the normal stress difference. Once the relaxation modulus is obtained, shear viscosity (*η*) is calculated by integrating the shear stress relaxation modulus over time using the last GK formula

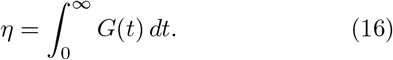

To mitigate noise in the relaxation modulus observed in protein condensate simulations^31,69^, we adopt a specific strategy for viscosity estimation. While *G*(*t*) is smooth at short timescales so that it can be numerically integrated, at longer timescales *G*(*t*) is fitted to a series of Maxwell modes (*G*_*i*_ exp(-*t/τ*_i_)) equidistant in logarithmic time, and then analytically integrated. Thus, viscosity is calculated by combining two terms as

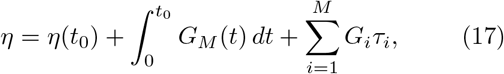

where *η*(*t*_0_) represents the term computed at short times, 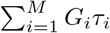 corresponds to the part evaluated via the Maxwell modes fit at long timescales, and *t*_0_ denotes the time after which all intramolecular oscillations of *G*(*t*) have decayed and the function becomes strictly positive and decays monotonously.

### F. Deviation from the ideal line in interpolations

For cases where simulated saturation concentration have been plotted against experimental values (Fig. 5 in the main text and Fig. B in SI, the following formula has been used to calculate the deviation of simulated results from experimental ones

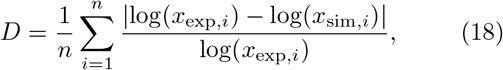

where log represents the decimal logarithm.

For cases where simulated viscosity have been plotted against experimental values (Fig. 6) the following formula has been used to calculate the deviation of simulated results from experimental ones

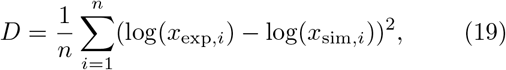

where log represents the decimal logarithm.

## VI. SUPPORTING INFORMATION

**TABLE A. Amino acid sequences of the different hnRNPA1-LCD variants**. The amino acid sequences of the hnRNPA1-LCD variants utilized in this investigation. In the first column, we have listed the names of the variants. The second column displays their amino acid sequences, while the third column indicates the number and type of changes compared to the wild-type sequence. **FIG. A. Relative interaction strength values for the HPS (A), HPS-cation-** *π* **(B), HPS-Urry (C), CALVADOS2 (D), Mpipi (E), and Mpipi-Recharged (F) models**. The values have been normalized by the highest interaction value of each model. **FIG. B. Simulated vs. experimental saturation concentration**^78^ **for the different variants and for the models CALVADOS2 (A), Mpipi (B) and Mpipi-Recharged (C)**. The Pearson correlation coefficient (r), the slope (m) and the root mean square deviation from the experimental values (D) are displayed for each set of modelling data. The black lines indicate a perfect match between experimental and computational values, while the red dashed lines depict the linear regression for each set of data. **FIG. C. Contact maps of potential energy interaction for the A1-LCD (WT+NLS) sequence** predicted by the HPS (A), HPS-cation-*π* (B), HPS-Urry (C), CALVADOS2 (D), Mpipi (E), and Mpipi-Recharged (F) models at T = 0.95 T_*c*_ (where T_*c*_ refers to the critical temperature of the WT+NLS sequence of each model) and at the condensate equilibrium density corresponding to such temperature. The associated energy corresponding to a given specific interaction is depicted by the side bar. The associated energy corresponding to a given specific interaction is depicted by the side bar. Details on these calculations are provided in the Methods Section. **FIG. D. Predominant intermolecular interactions contributing to LLPS of A1-LCD (WT+NLS) protein** as predicted by the HPS (A), HPS-cation-*π* (B), HPS-Urry (C), CALVADOS2 (D), and Mpipi (E) models. The presented contribution by each residue-residue pair has been normalised by the highest contact pair. Also, normalisation by the pair residue-residue abundance across the sequence has been applied. **SUPPORTING MOVIE A. Simulation movie of a representative trajectory of A1-LCD using the Direct Coexistence method to compute the phase diagram. SUPPORTING MOVIE B. Simulation movie of a representative trajectory of A1-LCD performed for the calculation of the condensate viscosity**.

## VII. ACKNOWLEDGEMENTS

A. F. acknowledges funding from the Ramon y Cajal fellowship (RYC2021-030937-I) and Spanish National Grant (PID2022-136919NA-C33) I. S.-B. acknowledges funding from the Derek Brewer scholarship of Emmanuel College and EPSRC Doctoral Training Programme studentship, number EP/T517847/1. R.C.-G. acknowledges funding from the European Research Council (ERC) under the European Union Horizon 2020 research and innovation programme (grant agreement 803326). A. T. is funded by European Research Council (ERC) under the European Union Horizon 2020 research and innovation programme (grant agreement 803326) and Ramon y Cajal fellowship (RYC2021-030937-I). J. R. E. also acknowledges funding from the Roger Ekins Research Fellowship of Emmanuel College, the Ramon y Cajal fellowship (RYC2021-030937-I) and the Spanish National Agency for Research (PID2022-136919NA-C33). A.R also acknowledges funding from PID2023-147156NB-I00 of the Spanish Ministry for Science, Innovation and Universities. This work has been performed using resources provided by the Cambridge Tier-2 system operated by the University of Cambridge Research Computing Service (http://www.hpc.cam.ac.uk) funded by EPSRC Tier-2 capital grant EP/P020259/1. The authors thankfully acknowledge RES computational resources provided by Mare Nostrum 5 through the activity 2024-3-0001.

## VIII. DATA AVAILABILITY STATEMENT

We provide Supplementary Material and a GitHub Repository^137^ that contains the LAMMPS input scripts for every studied model as well as the configuration file for the considered A1-LCD mutants. We also provide sampling videos of representative trajectories concerning the direct coexistence method, viscosity and different conformations of a protein.

## Notes

### Competing Interest Statement

The authors have declared no competing interest.

### Summary of Updates

We have ammended some typos, added some figures and include a new model in our analysis.

## REFERENCES

1 A. A. Hyman and K. Simons, Science 337, 1047 (2012).

2 A. A. Hyman, C. A. Weber, and F. Jülicher, Annual review of cell and developmental biology 30, 39 (2014).

3 C. P. Brangwynne, C. R. Eckmann, D. S. Courson, A. Rybarska, C. Hoege, J. Gharakhani, F. Jülicher, and A. A. Hyman, Science 324, 1729 (2009).

4 X. Su, J. A. Ditlev, E. Hui, W. Xing, S. Banjade, J. Okrut, D. S. King, J. Taunton, M. K. Rosen, and R. D. Vale, Science 352, 595 (2016).

5 Q. Su, S. Mehta, and J. Zhang, Molecular cell 81, 4137 (2021).

6 Q. Xiao, C. K. McAtee, and X. Su, Nature Reviews Immunology 22, 188 (2022).

7 S. Boeynaems, S. Alberti, N. L. Fawzi, T. Mittag, M. Polymenidou, F. Rousseau, J. Schymkowitz, J. Shorter, B. Wolozin, L. Van Den Bosch, et al., Trends in cell biology 28, 420 (2018).

8 H. Zhou, Z. Song, S. Zhong, L. Zuo, Z. Qi, L.-J. Qu, and L. Lai, Angewandte Chemie 131, 4912 (2019).

9 J. A. Lemkul and D. R. Bevan, The Journal of Physical Chemistry B 114, 1652 (2010).

10 A. Musacchio, The EMBO journal 41, e109952 (2022).

11 A. R. Strom, A. V. Emelyanov, M. Mir, D. V. Fyodorov, X. Darzacq, and G. H. Karpen, Nature 547, 241 (2017).

12 G. J. Narlikar, Journal of Biosciences 45, 5 (2020).

13 D. Deviri and S. A. Safran, Proceedings of the National Academy of Sciences 118, e2100099118 (2021).

14 J. A. Riback and C. P. Brangwynne, Science 367, 364 (2020).

15 A. Klosin, F. Oltsch, T. Harmon, A. Honigmann, F. Jülicher, A. A. Hyman, and C. Zechner, Science 367, 464 (2020).

16 J. Sheu-Gruttadauria and I. J. MacRae, Cell 173, 946 (2018).

17 J. J. Bouchard, J. H. Otero, D. C. Scott, E. Szulc, E. W. Martin, N. Sabri, D. Granata, M. R. Marzahn, K. Lindorff-Larsen, X. Salvatella, et al., Molecular cell 72, 19 (2018).

18 S. Qamar, G. Wang, S. J. Randle, F. S. Ruggeri, J. A. Varela, J. Q. Lin, E. C. Phillips, A. Miyashita, D. Williams, F. Ströhl, et al., Cell 173, 720 (2018).

19 A. C. Murthy, G. L. Dignon, Y. Kan, G. H. Zerze, S. H. Parekh, J. Mittal, and N. L. Fawzi, Nature structural & molecular biology 26, 637 (2019).

20 J. L. Carey and L. Guo, Frontiers in molecular biosciences 9, 826719 (2022).

21 H.-R. Li, W.-C. Chiang, P.-C. Chou, W.-J. Wang, and J.-r. Huang, Journal of Biological Chemistry 293, 6090 (2018).

22 L. McGurk, E. Gomes, L. Guo, J. Mojsilovic-Petrovic, V. Tran, R. G. Kalb, J. Shorter, and N. M. Bonini, Molecular cell 71, 703 (2018).

23 E. W. Martin, F. E. Thomasen, N. M. Milkovic, M. J. Cuneo, C. R. Grace, A. Nourse, K. Lindorff-Larsen, and T. Mittag, Nucleic acids research 49, 2931 (2021).

24 A. Molliex, J. Temirov, J. Lee, M. Coughlin, A. P. Kanagaraj, H. J. Kim, T. Mittag, and J. P. Taylor, Cell 163, 123 (2015).

25 L. Guo and J. Shorter, Molecular cell 60, 189 (2015).

26 X. Wang, J. C. Schwartz, and T. R. Cech, Nucleic acids research 43, 7535 (2015).

27 M. Farag, S. R. Cohen, W. M. Borcherds, A. Bremer, T. Mittag, and R. V. Pappu, Nature communications 13, 7722 (2022).

28 W. Borcherds, A. Bremer, M. B. Borgia, and T. Mittag, Current opinion in structural biology 67, 41 (2021).

29 M. P. Hughes, M. R. Sawaya, D. R. Boyer, L. Goldschmidt, J. A. Rodriguez, D. Cascio, L. Chong, T. Gonen, and D. S. Eisenberg, Science 359, 698 (2018).

30 S. Ray, N. Singh, R. Kumar, K. Patel, S. Pandey, D. Datta, J. Mahato, R. Panigrahi, A. Navalkar, S. Mehra, et al., Nature chemistry 12, 705 (2020).

31 A. R. Tejedor, I. Sanchez-Burgos, M. Estevez-Espinosa, A. Garaizar, R. Collepardo-Guevara, J. Ramirez, and J. R. Espinosa, Nature communications 13, 1 (2022).

32 P. P. Gopal, J. J. Nirschl, E. Klinman, and E. L. Holzbaur, Proceedings of the National Academy of Sciences 114, E2466 (2017).

33 P. St George-Hyslop, J. Q. Lin, A. Miyashita, E. C. Phillips, S. Qamar, S. J. Randle, and G. Wang, Brain research 1693, 11 (2018).

34 A. Patel, H. O. Lee, L. Jawerth, S. Maharana, M. Jahnel, M. Y. Hein, S. Stoynov, J. Mahamid, S. Saha, T. M. Franzmann, et al., Cell 162, 1066 (2015).

35 I. R. Mackenzie, A. M. Nicholson, M. Sarkar, J. Messing, M. D. Purice, C. Pottier, K. Annu, M. Baker, R. B. Perkerson, A. Kurti, et al., Neuron 95, 808 (2017).

36 Y. Lu, L. Lim, and J. Song, Scientific Reports 7, 1 (2017).

37 S. Ambadipudi, J. Biernat, D. Riedel, E. Mandelkow, and M. Zweckstetter, Nature communications 8, 1 (2017).

38 I. Bishof, E. B. Dammer, D. M. Duong, S. R. Kundinger, M. Gearing, J. J. Lah, A. I. Levey, and N. T. Seyfried, Journal of Biological Chemistry 293, 11047 (2018).

39 S. Spannl, M. Tereshchenko, G. J. Mastromarco, S. J. Ihn, and H. O. Lee, Traffic 20, 890 (2019).

40 S. Mehta and J. Zhang, Nature Reviews Cancer 22, 239 (2022).

41 G. Tesei, T. K. Schulze, R. Crehuet, and K. Lindorff-Larsen, Proceedings of the National Academy of Sciences 118, e2111696118 (2021).

42 B. Sza-la-Mendyk, T. M. Phan, P. Mohanty, and J. Mittal, Current Opinion in Chemical Biology 75, 102333 (2023).

43 G. L. Dignon, W. Zheng, Y. C. Kim, R. B. Best, and J. Mittal, PLoS computational biology 14, e1005941 (2018).

44 M. Paloni, R. Bailly, L. Ciandrini, and A. Barducci, The Journal of Physical Chemistry B 124, 9009 (2020).

45 W. Zheng, G. L. Dignon, N. Jovic, X. Xu, R. M. Regy, N. L. Fawzi, Y. C. Kim, R. B. Best, and J. Mittal, The Journal of Physical Chemistry B 124, 11671 (2020).

46 J. Sponer, M. Krepl, P. Banas, P. Kuhrova, M. Zgarbova, P. Jurecka, M. Havrila, and M. Otyepka, Wiley Interdisciplinary Reviews: RNA 8, e1405 (2017).

47 S. Torrino, W. Oldham, A. R. Tejedor, I. Sanchez-Burgos, N. Rachedi, K. Fraissard, C. Chauvet, C. Sbai, B. P. O’Hara, S. Abelanet, et al., “Mechano-dependent sorbitol accumulation supports biomolecular condensate,” (2023).

48 A. Garaizar, T. Higginbotham, I. Sanchez-Burgos, A. R. Tejedor, E. Sanz, and J. R. Espinosa, The Journal of Chemical Physics 157 (2022).

49 J. R. Espinosa, A. Garaizar, C. Vega, D. Frenkel, and R. Collepardo-Guevara, The Journal of chemical physics 150, 224510 (2019).

50 V. Nguemaha and H.-X. Zhou, Scientific reports 8, 1 (2018).

51 I. Sanchez-Burgos, J. R. Espinosa, J. A. Joseph, and R. Collepardo-Guevara, Biomolecules 11, 278 (2021).

52 G. L. Dignon, W. Zheng, Y. C. Kim, R. B. Best, and J. Mittal, PLoS computational biology 14, e1005941 (2018).

53 J. A. Joseph, A. Reinhardt, A. Aguirre, P. Y. Chew, K. O. Russell, J. R. Espinosa, A. Garaizar, and R. Collepardo-Guevara, Nature Computational Science 1, 732 (2021).

54 R. M. Regy, J. Thompson, Y. C. Kim, and J. Mittal, Protein Science 30, 1371 (2021).

55 G. Tesei and K. Lindorff-Larsen, Open Research Europe 2, 94 (2023).

56 S. Das, Y.-H. Lin, R. M. Vernon, J. D. Forman-Kay, and H. S. Chan, Proceedings of the National Academy of Sciences 117, 28795 (2020).

57 J. M. Lotthammer, G. M. Ginell, D. Griffith, R. Emenecker, and A. S. Holehouse, Biophysical Journal 123, 43a (2024).

58 T. Dannenhoffer-Lafage and R. B. Best, The Journal of Physical Chemistry B 125, 4046 (2021).

59 A. Garaizar and J. R. Espinosa, The Journal of Chemical Physics 155, 125103 (2021).

60 A. Statt, H. Casademunt, C. P. Brangwynne, and A. Z. Panagiotopoulos, The Journal of Chemical Physics 152, 075101 (2020).

61 A. R. Tejedor, A. Aguirre Gonzalez, M. J. Maristany, P. Y. Chew, K. Rusell, J. Ramirez, J. R. Espinosa, and R. Collepardo-Guevara, bioRxiv (2024), 10.1101/2024.07.26.605370.

62 A. C. Murthy, G. L. Dignon, Y. Kan, G. H. Zerze, S. H. Parekh, J. Mittal, and N. L. Fawzi, Nature structural & molecular biology 26, 637 (2019).

63 G. L. Dignon, R. B. Best, and J. Mittal, Annual Review of Physical Chemistry 71, 53 (2020).

64 G. L. Dignon, W. Zheng, and J. Mittal, Current opinion in chemical engineering 23, 92 (2019).

65 B. S. Schuster, G. L. Dignon, W. S. Tang, F. M. Kelley, A. K. Ranganath, C. N. Jahnke, A. G. Simpkins, R. M. Regy, D. A. Hammer, M. C. Good, et al., Proceedings of the National Academy of Sciences 117, 11421 (2020).

66 V. H. Ryan, G. L. Dignon, G. H. Zerze, C. V. Chabata, R. Silva, A. E. Conicella, J. Amaya, K. A. Burke, J. Mittal, and N. L. Fawzi, Molecular cell 69, 465 (2018).

67 A. Abyzov, M. Blackledge, and M. Zweckstetter, Chemical Reviews 122, 6719 (2022).

68 M. J. Maristany, A. A. Gonzalez, J. R. Espinosa, J. Huertas, R. Collepardo-Guevara, and J. A. Joseph, bioRxiv, 2023 (2023).

69 A. R. Tejedor, A. Garaizar, J. Ramírez, and J. R. Espinosa, Biophysical Journal 120, 5169 (2021).

70 I. Sanchez-Burgos, J. R. Espinosa, J. A. Joseph, and R. Collepardo-Guevara, PLoS computational biology 18, e1009810 (2022).

71 R. Funari, N. Bhalla, and L. Gentile, ACS Measurement Science Au 2, 547 (2022).

72 X. Huang and R. Powers, Journal of the American Chemical Society 123, 3834 (2001).

73 K. Araki, N. Yagi, R. Nakatani, H. Sekiguchi, M. So, H. Yagi, N. Ohta, Y. Nagai, Y. Goto, and H. Mochizuki, Scientific reports 6, 30473 (2016).

74 J. A. Riback, M. A. Bowman, A. M. Zmyslowski, C. R. Knoverek, J. M. Jumper, J. R. Hinshaw, E. B. Kaye, K. F. Freed, P. L. Clark, and T. R. Sosnick, Science 358, 238 (2017).

75 G.-N. W. Gomes, M. Krzeminski, A. Namini, E. W. Martin, T. Mittag, T. Head-Gordon, J. D. Forman-Kay, and C. C. Gradinaru, Journal of the American Chemical Society 142, 15697 (2020).

76 G. L. Dignon, W. Zheng, R. B. Best, Y. C. Kim, and J. Mittal, Proceedings of the National Academy of Sciences 115, 9929 (2018).

77 A. Bremer, M. Farag, W. M. Borcherds, I. Peran, E. W. Martin, R. V. Pappu, and T. Mittag, Nature Chemistry 14, 196 (2022).

78 I. Alshareedah, W. M. Borcherds, S. R. Cohen, A. Singh, A. E. Posey, M. Farag, A. Bremer, G. W. Strout, D. T. Tomares, R. V. Pappu, et al., Nature Physics, 1 (2024).

79 J. A. Joseph, A. Reinhardt, A. Aguirre, P. Y. Chew, K. O. Russell, J. R. Espinosa, A. Garaizar, and R. Collepardo-Guevara, Nature Computational Science 1, 732 (2021).

80 P. Li, S. Banjade, H.-C. Cheng, S. Kim, B. Chen, L. Guo, M. Llaguno, J. V. Hollingsworth, D. S. King, S. F. Banani, et al., Nature 483, 336 (2012).

81 S. Boeynaems, S. Alberti, N. L. Fawzi, T. Mittag, M. Polymenidou, F. Rousseau, J. Schymkowitz, J. Shorter, B. Wolozin, L. Van Den Bosch, et al., Trends in cell biology 28, 420 (2018).

82 R. M. Vernon, P. A. Chong, B. Tsang, T. H. Kim, A. Bah, P. Farber, H. Lin, and J. D. Forman-Kay, elife 7, e31486 (2018).

83 E. W. Martin, A. S. Holehouse, I. Peran, M. Farag, J. J. Incicco, A. Bremer, C. R. Grace, A. Soranno, R. V. Pappu, and T. Mittag, Science 367, 694 (2020).

84 A. Ladd and L. Woodcock, Chemical Physics Letters 51, 155 (1977).

85 J. R. Espinosa, J. A. Joseph, I. Sanchez-Burgos, A. Garaizar, D. Frenkel, and R. Collepardo-Guevara, Proceedings of the National Academy of Sciences 117, 13238 (2020).

86 L. H. Kapcha and P. J. Rossky, Journal of molecular biology 426, 484 (2014).

87 J. Wessén, S. Das, T. Pal, and H. S. Chan, The Journal of Physical Chemistry B 126, 9222 (2022).

88 D. W. Urry, D. C. Gowda, T. M. Parker, C.-H. Luan, M. C. Reid, C. M. Harris, A. Pattanaik, and R. D. Harris, Biopolymers: Original Research on Biomolecules 32, 1243 (1992).

89 A. Feito, I. Sanchez-Burgos, A. Rey, R. Collepardo-Guevara, J. R. Espinosa, and A. R. Tejedor, Molecular Physics, e2425757 (2024).

90 A. Ladd and L. Woodcock, Chemical Physics Letters 51, 155 (1977).

91 T. Pal, J. Wessén, S. Das, and H. S. Chan, The Journal of Physical Chemistry Letters 15, 8248 (2024).

92 S. Alberti and D. Dormann, Annu. Rev. Genet 53, 171 (2019).

93 M.-T. Wei, S. Elbaum-Garfinkle, A. S. Holehouse, C. C.-H. Chen, M. Feric, C. B. Arnold, R. D. Priestley, R. V. Pappu, and C. P. Brangwynne, Nature Chemistry 9, 1118 (2017).

94 A. Wang, A. E. Conicella, H. B. Schmidt, E. W. Martin, S. N. Rhoads, A. N. Reeb, A. Nourse, D. Ramirez Montero, V. H. Ryan, R. Rohatgi, et al., The EMBO journal 37, e97452 (2018).

95 S. Maharana, J. Wang, D. K. Papadopoulos, D. Richter, A. Pozniakovsky, I. Poser, M. Bickle, S. Rizk, J. Guillen-Boixet, T. M. Franzmann, M. Jahnel, L. Marrone, Y.-T. Chang, J. Sterneckert, P. Tomancak, A. A. Hyman, and S. Alberti, Science 360, 918 (2018).

96 J. Wang, J.-M. Choi, A. S. Holehouse, H. O. Lee, X. Zhang, M. Jahnel, S. Maharana, R. Lemaitre, A. Pozniakovsky, D. Drechsel, et al., Cell 174, 688 (2018).

97 C. N. Johnson, K. A. Sojitra, E. J. Sohn, A. K. Moreno-Romero, A. Baudin, X. Xu, J. Mittal, and D. S. Libich, Journal of the American Chemical Society 146, 8071 (2024).

98 W. M. Babinchak, R. Haider, B. K. Dumm, P. Sarkar, K. Surewicz, J.-K. Choi, and W. K. Surewicz, Journal of Biological Chemistry 294, 6306 (2019).

99 M. Hofweber, S. Hutten, B. Bourgeois, E. Spreitzer, A. Niedner-Boblenz, M. Schifferer, M.-D. Ruepp, M. Simons, D. Niessing, T. Madl, et al., Cell 173, 706 (2018).

100 G. Krainer, T. J. Welsh, J. A. Joseph, J. R. Espinosa, S. Wittmann, E. de Csillery, A. Sridhar, Z. Toprakcioglu, G. Gudiškytė, M. A. Czekalska, et al., Nature Communications 12, 1 (2021).

101 S. Alberti and A. A. Hyman, Nature reviews Molecular cell biology 22, 196 (2021).

102 B. Portz, B. L. Lee, and J. Shorter, Trends in biochemical sciences 46, 550 (2021).

103 L. Jawerth, E. Fischer-Friedrich, S. Saha, J. Wang, T. Franzmann, X. Zhang, J. Sachweh, M. Ruer, M. Ijavi, S. Saha, et al., Science 370, 1317 (2020).

104 S. Chatterjee, Y. Kan, M. Brzezinski, K. Koynov, R. M. Regy, A. C. Murthy, K. A. Burke, J. J. Michels, J. Mittal, N. L. Fawzi, et al., Advanced Science 9, 2104247 (2022).

105 E. L. Guenther, Q. Cao, H. Trinh, J. Lu, M. R. Sawaya, D. Cascio, D. R. Boyer, J. A. Rodriguez, M. P. Hughes, and D. S. Eisenberg, Nature structural & molecular biology 25, 463 (2018).

106 M. Linsenmeier, L. Faltova, C. Morelli, U. Capasso Palmiero, C. Seiffert, A. M. Küffner, D. Pinotsi, J. Zhou, R. Mezzenga, and P. Arosio, Nature chemistry 15, 1340 (2023).

107 R. S. Fisher and S. Elbaum-Garfinkle, Nature communications 11, 1 (2020).

108 D. Sundaravadivelu Devarajan, J. Wang, B. Sza-la-Mendyk, S. Rekhi, A. Nikoubashman, Y. C. Kim, and J. Mittal, Nature Communications 15, 1912 (2024).

109 S. Boeynaems, A. S. Holehouse, V. Weinhardt, D. Kovacs, J. Van Lindt, C. Larabell, L. Van Den Bosch, R. Das, P. S. Tompa, R. V. Pappu, et al., Proceedings of the National Academy of Sciences 116, 7889 (2019).

110 J. Ramirez, S. K. Sukumaran, B. Vorselaars, and A. E. Likhtman, The Journal of chemical physics 133, 154103 (2010).

111 A. R. Tejedor, J. R. Tejedor, and J. Ramírez, The Journal of Chemical Physics 157 (2022).

112 A. R. Tejedor, R. Collepardo-Guevara, J. Ramirez, and J. R. Espinosa, The Journal of Physical Chemistry B 127, 4441 (2023).

113 H. Deng, K. Gao, and J. Jankovic, Nature Reviews Neurology 10, 337 (2014).

114 S. Wegmann, B. Eftekharzadeh, K. Tepper, K. M. Zoltowska, R. E. Bennett, S. Dujardin, P. R. Laskowski, D. MacKenzie, T. Kamath, C. Commins, et al., The EMBO journal 37, e98049 (2018).

115 S. K. Rai, A. Savastano, P. Singh, S. Mukhopadhyay, and M. Zweckstetter, Protein Science 30, 1294 (2021).

116 N. Galvanetto, M. T. Ivanović, A. Chowdhury, A. Sottini, M. F. Nüesch, D. Nettels, R. B. Best, and B. Schuler, Nature 619, 876 (2023).

117 J.-M. Choi and R. V. Pappu, Biophysical Journal 118, 492a (2020).

118 U. B. Choi, H. Sanabria, T. Smirnova, M. E. Bowen, and K. R. Weninger, Biomolecules 9, 114 (2019).

119 H.-X. Zhou, D. Kota, S. Qin, and R. Prasad, Chemical Reviews (2024).

120 R. M. Regy, G. L. Dignon, W. Zheng, Y. C. Kim, and J. Mittal, Nucleic acids research 48, 12593 (2020).

121 H. S. Ashbaugh and H. W. Hatch, Journal of the American Chemical Society 130, 9536 (2008).

122 H. A. Lorentz, Annals of physics 248, 127 (1881).

123 D. Berthelot, Compt. Rendus 126, 15 (1898).

124 S. Plimpton, Journal of computational physics 117, 1 (1995).

125 A. P. Thompson, H. M. Aktulga, R. Berger, D. S. Bolintineanu, W. M. Brown, P. S. Crozier, P. J. in ‘t Veld, A. Kohlmeyer, S. G. Moore, T. D. Nguyen, R. Shan, M. J. Stevens, J. Tranchida, C. Trott, and S. J. Plimpton, Comp. Phys. Comm. 271, 108171 (2022).

126 T. Schneider and E. Stoll, Physical Review B 17, 1302 (1978).

127 R. García Fernández, J. L. Abascal, and C. Vega, The Journal of chemical physics 124 (2006).

128 F. J. Blas, L. G. MacDowell, E. de Miguel, and G. Jackson, The Journal of chemical physics 129 (2008).

129 J. R. Espinosa, E. Sanz, C. Valeriani, and C. Vega, The Journal of chemical physics 139 (2013).

130 J. S. Rowlinson and B. Widom, Molecular theory of capillarity (Courier Corporation, 2013).

131 M. Rubinstein and R. H. Colby, Polymer physics (Oxford university press, 2003).

132 A. E. Likhtman, S. K. Sukumaran, and J. Ramirez, Macro-molecules 40, 6748 (2007).

133 A. R. Tejedor, J. R. Tejedor, and J. Ramírez, The Journal of Chemical Physics 157 (2022).

134 J. Ramírez, S. K. Sukumaran, B. Vorselaars, and A. E. Likhtman, The Journal of chemical physics 133 (2010).

135 D. Bagheriasl, P. J. Carreau, B. Riedl, C. Dubois, and W. Y. Hamad, Cellulose 23, 1885 (2016).

136 R. S. Fisher and S. Elbaum-Garfinkle, Biophysical Journal 121, 147a (2022).

137 The GitHub repository can be accessed in this URL: https://github.com/Reshiiiii/hnRNPA1_Data_Scripts.

