## Supplementary Material for "Benchmarking residue-resolution protein coarse-grained models for simulations of biomolecular condensates"

Ignacio Sanchez-Burgos

*Maxwell Centre, Cavendish Laboratory, Department of Physics, University of Cambridge, J J Thomson Avenue, Cambridge CB3 0HE, United Kingdom.*

Rosana Collepardo-Guevara

*Yusuf Hamied Department of Chemistry, University of Cambridge, Lensfield Road, Cambridge CB2 1EW, UK and Department of Genetics, University of Cambridge, Cambridge CB2 3EH, UK*

Andrés R. Tejedor†

*Department of Physical-Chemistry Universidad Complutense de Madrid Av. Complutense s/n, Madrid 28040, Spain and Yusuf Hamied Department of Chemistry, University of Cambridge, Lensfield Road, Cambridge CB2 1EW, UK*

(Dated: December 19, 2024)

| Variant name | Protein sequence | Number and type of change |
|---|---|---|
| **WT+NLS** | GSMASASSSQRGRSGSGNFGGGRGGGFGGNDNFGRGGNFSGRGGFGG<br>SRGGGGYGGSGDGYNGFGNDGSNFGGGGSYNDFGNYNNQSSNFGPMK<br>GGNFGGRSSGPYGGGGQYFAKPRNQGGYGGSSSSSSYGSGRRF | |
| **allW** | GSMASASSSQRGRSGSGNWGGGRGGGWGGNDNWGRGGNWSGRGGWGG<br>SRGGGGWGGSGDGWNGWGNDGSNWGGGGSWNDWGNWNNQSSNWGPMK<br>GGNWGGRSSGSGGGGGQWWAKPRNQGGWGGSSSSSSWGSGRRW | 19W |
| **allY** | GSMASASSSQRGRSGSGNYGGGRGGGYGGNDNYGRGGNYSGRGGYGG<br>SRGGGGYGGSGDGYNGYGNDGSNYGGGGSYNDYGNYNNQSSNYGPMK<br>GGNYGGRSSGGSGGGGQYYAKPRNQGGYGGSSSSSSYGSGRRY | 19Y |
| **allF** | GSMASASSSQRGRSGSGNFGGGRGGGFGGNDNFGRGGNFSGRGGFGG<br>SRGGGGFGGSGDGFNGFGNDGSNFGGGGSFNDFGNFNNQSSNFGPMK<br>GGNFGGRSSGGSGGGGQFFAKPRNQGGFGGSSSSSSFGSGRRF | 19F |
| **W-** | GSMASASSSQRGRSGSGNSGGGRGGGWGGNDNWGRGGNSSGRGGWGG<br>SRGGGGWGGSGDGWNGWGNDGSNSGGGGSSNDWGNWNNQSSNWGPMK<br>GGNWGGRSSGGSGGGGQWSAKPRNQGGWGGSSSSSSSGSGRRW | 13W |
| **FtoW** | GSMASASSSQRGRSGSGNWGGGRGGGWGGNDNWGRGGNWSGRGGWGG<br>SRGGGGYGGSGDGYNGWGNDGSNWGGGGSYNDWGNYNNQSSNWGPMK<br>GGNWGGRSSGSGGGGGQYWAKPRNQGGYGGSSSSSSYGSGRRW | 12W 7Y |
| **YtoW** | GSMASASSSQRGRSGSGNFGGGRGGGFGGNDNFGRGGNFSGRGGFGG<br>SRGGGGWGGSGDGWNGFGNDGSNFGGGGSWNDFGNWNNQSSNFGPMK<br>GGNFGGRSSGSGGGGGQWFAKPRNQGGWGGSSSSSSWGSGRRF | 7W 12F |

**TABLE A:** The amino acid sequences of the hnRNPA1-LCD variants utilized in this investigation. In the first column, we have listed the names of the variants. The second column displays their amino acid sequences, while the third column indicates the number and type of changes compared to the wild-type sequence.


*

†

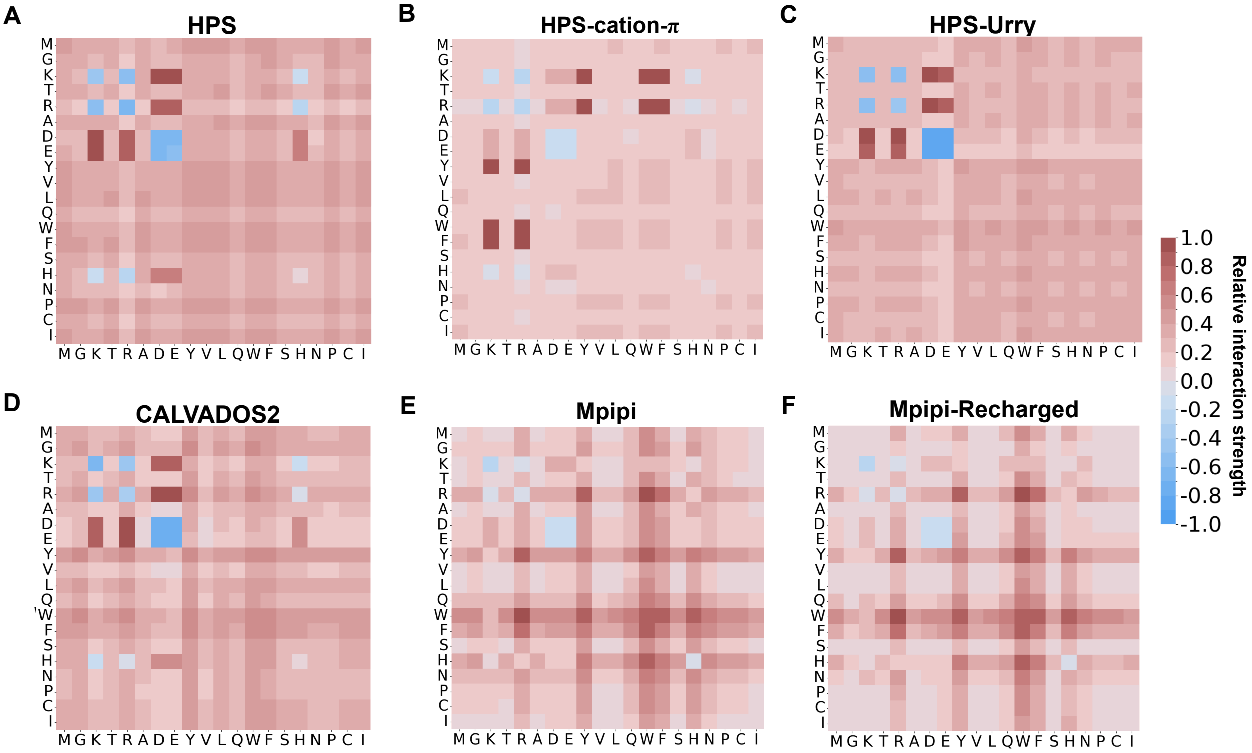


**FIG. A:** Predicted relative interaction strength values for the HPS (A), HPS-cation-$\pi$ (B), HPS-Urry (C), CALVADOS2 (D), Mpipi (E), and Mpipi-Recharged (F) models. The values have been normalized by the highest interaction value of each model.

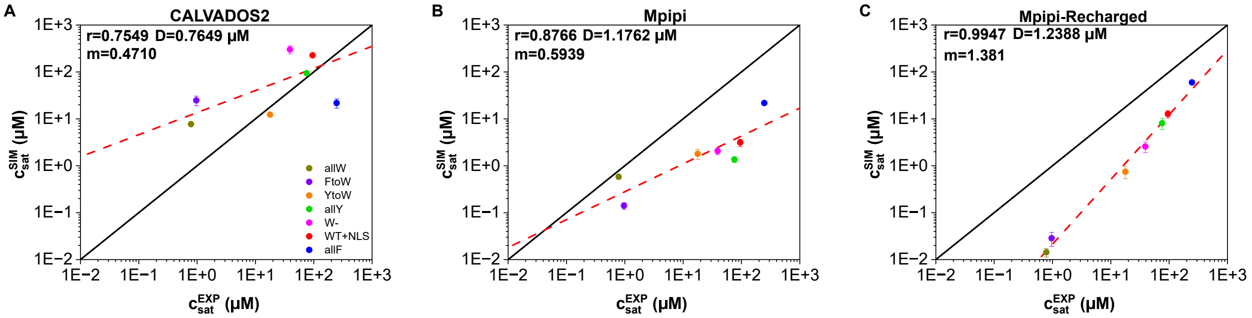


**FIG. B:** Simulated *vs.* experimental saturation concentration for the different variants and for the models CALVADOS2 (A), Mpipi (B) and Mpipi-Recharged (C). The Pearson correlation coefficient ($r$), the slope ($m$) and the root mean square deviation from the experimental values ($D$) are displayed for each set of modelling data. The black lines indicate a perfect match between experimental and computational values, while the red dashed lines depict the linear regression for each set of data.

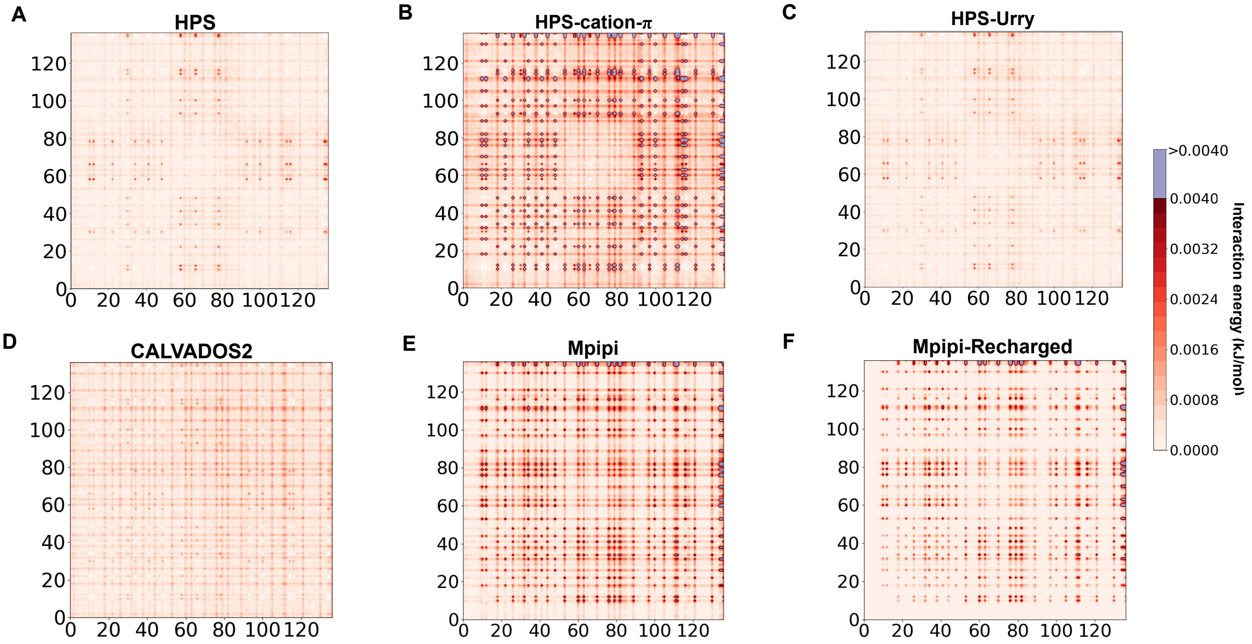


**FIG. C:** Contact maps of energy interaction for the A1-LCD (WT+NLS) sequence predicted by the HPS (A), HPS-cation–$\pi$ (B), HPS-Urry (C), CALVADOS2 (D), Mpipi (E), and Mpipi-Recharged (F) models at T = 0.95 $T_c$ (where $T_c$ refers to the critical temperature of the WT+NLS sequence of each model) at the condensate equilibrium density corresponding to such temperature. The associated energy corresponding to a given specific interaction is depicted by the side bar. Details on these calculations are provided in Section V D.

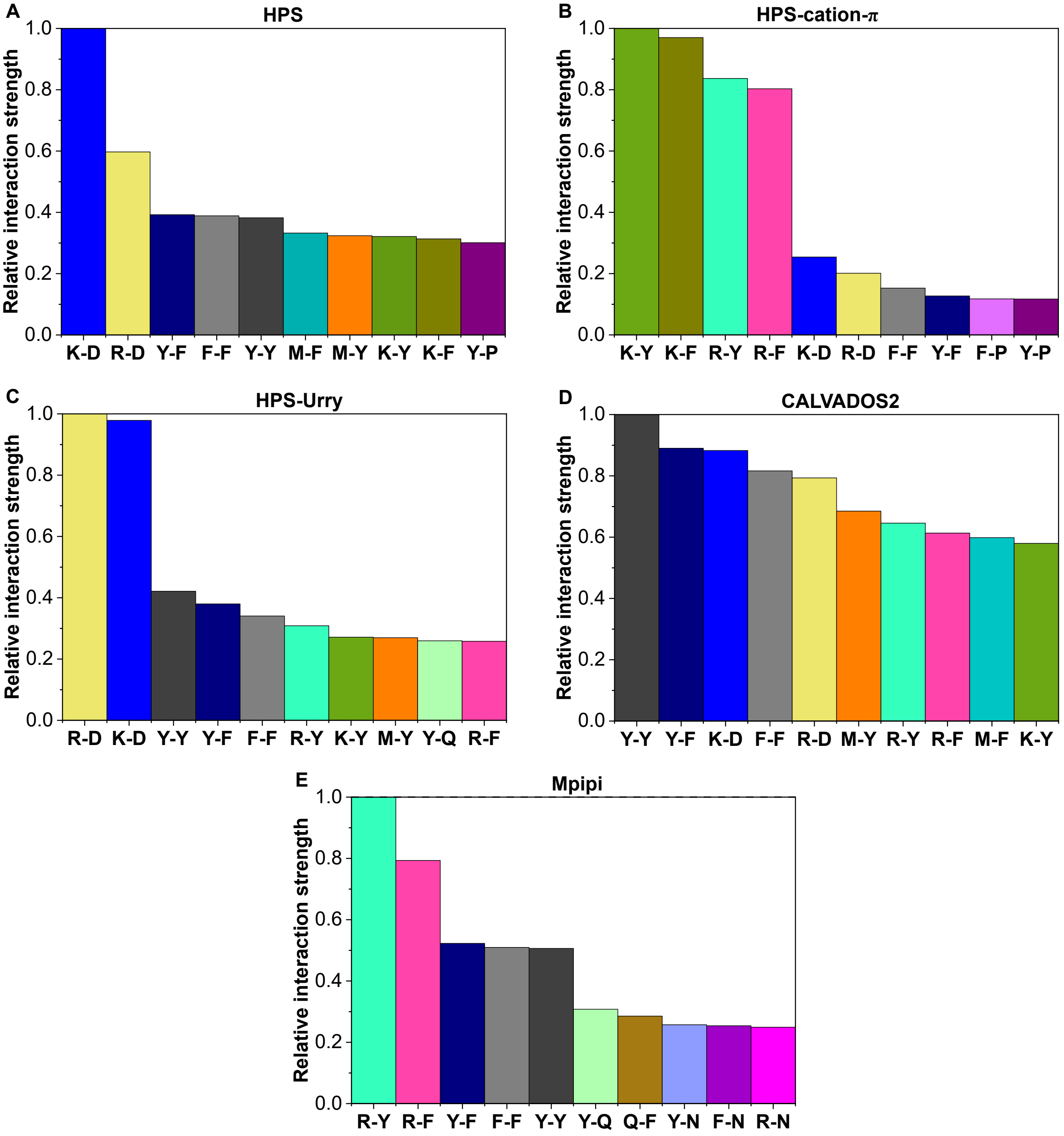


**FIG. D:** Predominant intermolecular interactions contributing to LLPS of A1-LCD (WT+NLS) protein as predicted by the HPS (A), HPS-cation–$\pi$ (B), HPS-Urry (C), CALVADOS2 (D), and Mpipi (E) models. The presented contribution by each residue-residue pair has been normalised by the highest contact pair. Also, normalisation by the pair residue-residue abundance across the sequence has been applied.
